# EZH2 inhibition in glioblastoma stem cells increases the expression of neuronal genes and the neuronal developmental regulators ZIC2, ZNF423 and MAFB

**DOI:** 10.1101/2021.11.22.469535

**Authors:** Bnar Abdul Kader, Rebecca Distefano, Katherine L. West, Adam G. West

## Abstract

Glioblastoma multiforme (GBM) is an aggressive brain cancer with a very poor prognosis. It has been shown that GBM stem cells within a GBM tumour have increased resistance to standard therapies, so new approaches are needed to increase the range of treatment options available. Here we use two GBM stem cell lines, representing the classical/pro-neural and mesenchymal GBM subtypes, to investigate the effects of three different EZH2 inhibitors on GBM stem cell survival and gene expression: EPZ6438, GSK343 and UNC1999. EZH2 is the catalytic component of the PRC2 chromatin repressor complex, which represses transcription through methylation of histone H3 at lysine 27. Both cell lines showed significantly reduced colony formation after 48-hour exposure to the inhibitors, indicating they were sensitive to all three EZH2 inhibitors. RNA-seq analysis revealed that all three EZH2 inhibitors led to increased expression of genes related to neurogenesis and/or neuronal structure in both GBM stem cell lines. Chromatin immunoprecipitation (ChIP-Seq) was used to identify potential direct targets of the histone methylation activity of EZH2 that might be driving the increase in neuronal gene expression. Three genes were identified as candidate regulatory targets common to both cell lines: MAFB, ZIC2 and ZNF423. These transcription factors all have known roles in regulating neurogenesis, brain development and/or neuronal function. Through analysis of three different EZH2 inhibitors and two GBM stem cell lines, this study demonstrates a common underlying mechanism for how inhibition of EZH2 activity reduces GBM stem cell proliferation and survival.

## Introduction

Glioblastoma multiforme (GBM), also known as grade IV astrocytoma, is the most prevalent and aggressive type of primary brain cancer, with an average survival duration from 15 to 30 months (Romani et al., 2018). Despite extensive knowledge of glioblastoma pathogenesis and a defined optimum treatment protocol, GBM remains incurable (Alifieris and Trafalis, 2015). Complete surgical removal of GBM is not achievable, due to the tumour’s infiltrating nature, so surgery is often coupled with radiotherapy and chemotherapy with Temozolomide (Alifieris and Trafalis, 2015).

GBMs can be classified into four main groups based on their pathology, genetic mutations and transcriptional profile: classical, mesenchymal, pro-neural and neural (Verhaak et al., 2010), and intra- and inter-tumour heterogenicity is one reason why development of targeted therapies has been challenging (Jin et al., 2017). Tumour complexity is also aggravated by the presence of glioblastoma cancer stem cells (GSCs), which have been linked to therapy resistance and poor prognosis due to their ability to self-renew and induce tumorigenesis (Singh et al., 2004, Bao et al., 2006). GSCs are mostly found in perivascular and hypoxic regions within the tumour, proliferating in a microenvironment characterised by poor delivery of oxygen and other nutrients (Wei et al., 2011, Zhu et al., 2011). As a result, drug penetration to these regions is also limited, explaining one of the basic means used by GSCs to escape therapy (Rosso et al., 2009, Seidel et al., 2010). Furthermore, cancer stem cells tend to migrate within the brain, supporting the infiltrating nature of the tumour and its ability to evade surgical removal (Hu et al., 2016, Krusche et al., 2016). Finally, they usually harbour genetic mutations that allow them to further resist treatments, supporting recurrent growth (Auffinger et al., 2015, Carruthers et al., 2015).

Epigenome manipulation is a key therapeutic area that is currently under investigation for many cancers to its essential role in the regulation of cell identity (Chammas et al., 2020). The interaction between chromatin regulatory proteins, long non-coding RNA, histones and non-histone proteins determines genome accessibility and gene expression. The primary regulators of gene silencing are Polycomb Repressive Complexes (PRC1 and PRC2) whose dysregulation has been observed in different types of cancer, including glioblastoma (Chang and Hung, 2012, Piunti and Pasini, 2011). PRC2 comprises SUZ12 (Suppressor of Zeste 12), EED (Embryonic Ectoderm Development), RBBP7 /RbAp46 (retinoblastoma-binding protein 7) and EZH1/-2 (Enhancer of Zeste Homolog 1 or 2) (Laugesen et al., 2019). Following its recruitment, PRC2 catalyses the tri-methylation of lysine 27 on histone H3 (H3K27me3) which enables the binding of PRC1 through the chromodomain of the CBX subunit. This, in turn, results in the monoubiquitylation of lysine 119 of histone H2A (H2AK119ub) by the RING1 E3 ubiquitin ligase of PRC1. The H3K27me3 mark is also recognised by the WD40-repeat domain of EED in PRC2, which causes a conformational change of EZH2, resulting in the activation of the methyltransferase activity of PRC2 and the methylation of neighbouring unmodified H3K27. This creates a feed-forward-loop that compacts the chromatin and causes repression of genes involved in the regulation of cell-cycle, self-renewal and cell differentiation (Yin et al., 2016, Laugesen et al., 2019).

EZH2 is a SAM (S-adenosyl-L-methionine) dependent methyltransferase and the main catalytic subunit of PRC2. While both EZH2 and EZH1 are able to catalyse mono- and di-methylation of H3K27, EZH2 alone can be fully activated by allosteric modulators and catalyse H3K27me3 (Lee et al., 2018). Consistently, *in vitro* studies have shown increased methyltransferase efficiency by EZH2 compared to EZH1, with EZH1 not being able to compensate for EZH2 loss of function (O’Carroll et al., 2001). Furthermore, while EZH1 is mostly expressed in differentiating cells, EZH2 is primarily associated with proliferating cells (Margueron et al., 2008). EZH2, and other PRC2 subunits, have been found overexpressed in different types of cancer, including breast, prostate, liver and GBM, where their upregulation has been linked to poor prognosis (Zhang et al., 2015). This is thought to be caused by PRC2-mediated silencing of tumour suppressor genes, including *TP53, PTEN, CDKN1A, CDKN2A, CDKN2B, RASSF1A and RARβ* (Beckedorff et al., 2013, Kazanets et al., 2016, Moison et al., 2014, Palakurthy et al., 2009, Paul et al., 2010, Schlesinger et al., 2007)

Beyond its role as methyltransferase in PRC2, EZH2 has also been reported to act on non-histone proteins in a PRC2-dependent and PRC2-independent way. For example, Kim *et al*. reported the phosphorylation of EZH2 by AKT, followed by the direct binding of EZH2 to STAT3, promoting tumour progression in glioblastoma and in GSCs (Kim et al., 2013), while Yi *et al*. observed that EZH2 overexpression was linked to the acceleration of ribosome function and altered translation control, resulting in cancer progression (Yi et al., 2021). Furthermore, interaction of EZH2 with SNAIL1 has also been observed, causing a decreased expression of E-cadherin and the subsequent promotion of Epithelial-to-Mesenchymal-Transition, resulting in the most aggressive GBM subgroup (mesenchymal) and with worst clinical outcome (Appolloni et al., 2015, Cao et al., 2008, Segerman et al., 2016).

Studies have already demonstrated that repression of EZH2 both *in vitro* and *in vivo* results in the reduction of cell proliferation and tumorigenesis, along with downregulation of cell cycle regulators and activation of apoptotic pathways (Fan et al., 2014, Suva et al., 2009, Zhang et al., 2015). Hence, due to its potential as a new therapeutic target, not only in GBM, several compounds have been synthesised and evaluated as EZH2 inhibitors.

One of the main classes of EZH2 inhibitor act as SAM-competitors, competitively binding to EZH2, decreasing its transmethylation ability. Current SAM-competitors being investigated include EPZ-6438, GSK-343 and UNC1999.

EPZ-6438 (Tazametostat) is an orally available molecule, with a higher affinity and selectivity for EZH2 compared to EZH1 and other histone methyltransferases (Knutson et al., 2014). Studies have shown its ability to inhibit H3K27me3 in a dose-dependent manner in lymphoma cell lines, while increasing percentage of cells in G1 phase and decreasing cell proliferation. Furthermore, induction of apoptotic cell death, increased medial survival and even complete eradication of tumours were observed on EZH2-Mutant Non-Hodgkin Lymphoma and medulloblastoma (Knutson et al., 2014, Zhang et al., 2015). A recent study on GBM U-87 cells, which targeted both EZH2 and P13K signalling through a combination drug therapy involving EPZ-6438, showed inhibition of GBM stemness, significant reduction of invasiveness, and a decrease in EMT markers (β-catenin, SNAIL3 and SMAD1) (Mishra et al., 2020). EPZ-6438 has been recently approved by the FDA for patients aged ≥ 16 with advanced epithelioid sarcoma (Duan et al., 2020), making it the first epigenetic drug approved for solid tumours. Furthermore, EPZ-6438 is currently under several phase I / II clinical trials for different lymphoma variants and several advanced solid tumours, including melanoma, lymphomas, urothelial carcinoma and metastatic prostate cancer (Gulati et al., 2018).

Similar results were also observed for GSK-343, a small molecule selective for EZH2 conceived in 2012 and still in the pre-clinical trial phase (Yu et al., 2017). Studies on ovarian, liver, colorectal cancers and glioblastoma showed decrease in colony formation in a dose-dependent manner and accumulation of G0/G1 cells (Amatangelo et al., 2013, Hsieh et al., 2016, Yu et al., 2017, Liu et al., 2015). Additional evidence showed a decrease in the levels of mesenchymal (N-cadherin, Vimentin, MMP2, MMP9, Snail and Slug) and pluripotency markers (Nestin, Sox2 and Oct-4), while stimulating an upregulation of tumour-suppressor genes commonly repressed by EZH2 (E-cadherin, *PTEN* and *P21*)(Fan et al., 2014, Liu et al., 2015, Xiong et al., 2020).

Finally, UNC1999, in contrast to EPZ-6438 and GSK-343, was developed as a dual inhibitor of EZH2 and EZH1. Similar to the two previously discussed drugs, studies on colon cancer cells, multiple myeloma and leukaemia have shown reduction of H3K27me3 in a dose dependent manner, induction of apoptosis, and downregulation of cell signalling genes (Alzrigat et al., 2017, Xu et al., 2015). A study of EZH2 inhibitors on GBM reported an increased specificity of UNC1999 due to its ability to inhibit H3K27me3, both *in vitro* and *in vivo*, without affecting the total level of H3 or EZH2, coupled with an observed decrease of GBM viable cell number and self-renewal capacity and increase of cells in the G1 phase (Grinshtein et al., 2016).

Overall, these studies highlight the anti-tumour properties of EPZ-6438, GSK-343 and UNC1999, and their ability to halt tumour cells proliferation through EZH2 inhibition. However, the exact mechanisms driving this inhibition is not clear. This, coupled with the variety of results obtained in glioblastoma treated cells highlight the need for a deeper understanding of the drivers behind the anti-tumour properties of these drugs. Here, we have investigated the transcriptomic and epigenetic changes following treatment with these three inhibitors in two different GBM cell lines, in order to identify common molecular mechanisms that underpin the cellular responses to these drugs.

## Results

### Inhibition of EZH2 reduces the proliferation of glioblastoma stem cell lines

Two GBM stem cell (GSC) lines were used to investigate the effect of treatment with EZH2 inhibitors, E2 and G7 (Gomez-Roman et al., 2017, Carruthers et al., 2015). RNA-seq analysis was used to investigate how similar these lines were to each other, and how they differed from normal neural stem cells (NSCs). E2 cells have higher expression of genes typically associated with the classical and pro-neural subtypes of GBM (Figure 1a) (Verhaak et al., 2010). This was supported by gene ontology analysis showing enrichment of categories associated with neurogenesis and neuronal function (Supplementary table 1) and gene set enrichment analysis (GSEA) showing increased expression of genes activated by hedgehog signalling (Supplementary table 2). Indeed, SHH and its receptor PTCH1 were both increased in E2 cells compared to G7. In contrast, G7 cells had higher expression of genes typically associated with the mesenchymal subtype of GBM (Figure 1a). GO analysis revealed enrichment in extracellular matrix genes (supplementary table 1), and GSEA indicated that the epithelial-mesenchymal-transition geneset was enriched in G7 compared to E2 cells (supplementary table 2). Variant analysis of RNA sequencing data revealed that the major cancer driver mutations in E2 cells include an activating mutation in EGFR (Ala289Val), a heterozygous deleterious mutation in TP53 (Arg273His), and biallelic loss of the CDKN2A locus (Supplementary table 5). Driver mutations in G7 cells include two deleterious heterozygous TP53 mutations (Arg248Gln and Arg282Trp). These analyses indicate that E2 and G7 cells represent different subtypes of GSCs.

**Figure 1.**
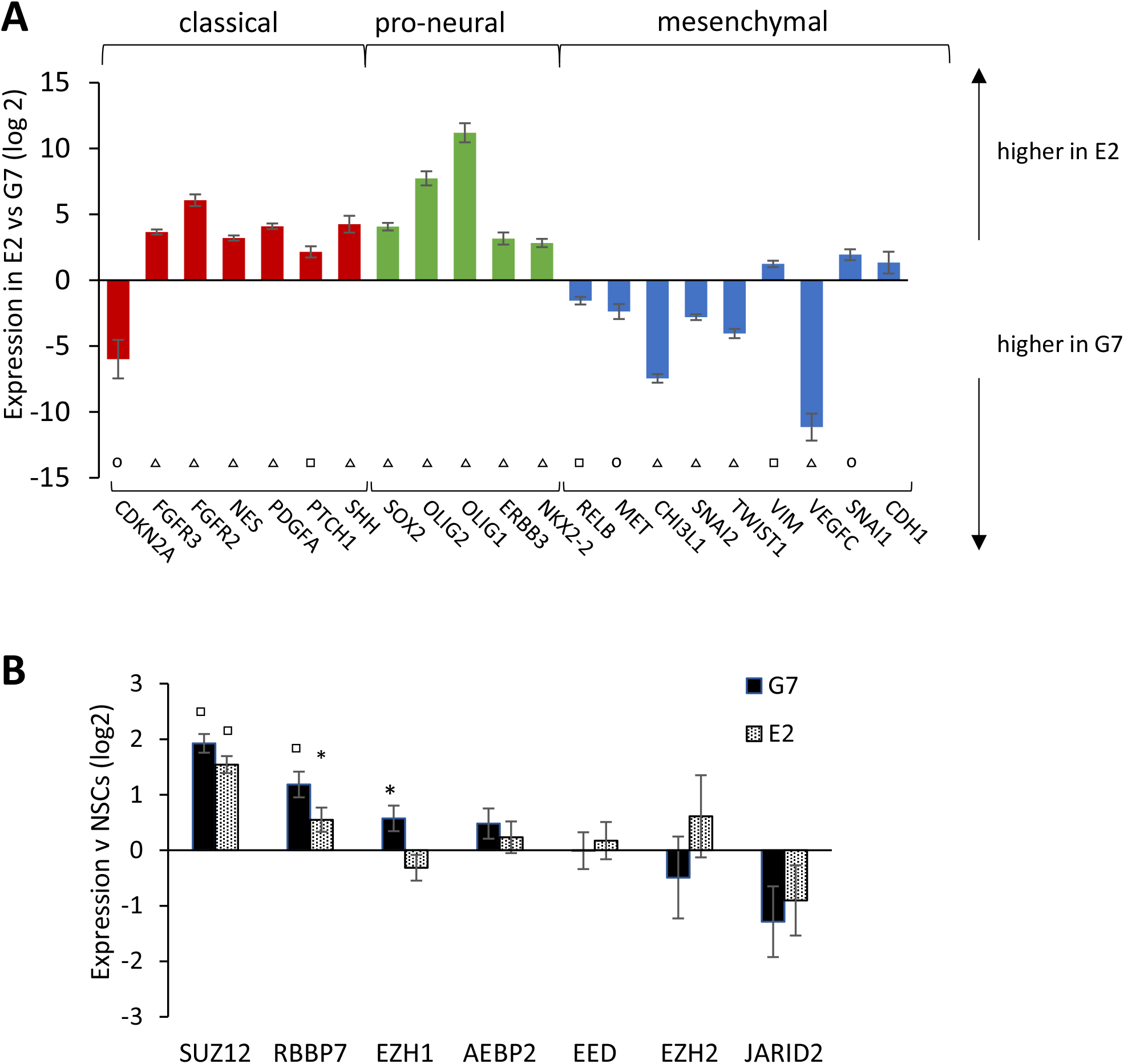
GBM cell lines overexpress PRC2 complex components. A) **Expression of genes representing GBM subtypes in E2 compared to G7 cells**. Genes were selected based on the analysis presented by Verhaak 2009. RNAseq data from E2 and G7 cells was analysed using DESEQ2, and the fold change in expression between E2 and G7 cells is plotted on a log2 scale for individual genes. O = p < 0.001, □ = p < 10^−5^, △ = p < 10^−10^ B) **Upregulation of key PRC2 genes in E2 and G7 cells compared to normal NSCs**. The relative expression of genes in the PRC2 complex was calculated using RNA-seq data from E2 and G7 cells, and from nine normal neural stem cell lines (Mack et al 2019). The difference in expression between E2 or G7 cells compared to the average expression in the normal NSCs is plotted as the mean +/- standard error on a log 2 scale. * = p □ < 0.05, = p < 10^−6^.

Increased expression of EZH2 and SUZ12 have been reported in gliomas compared to normal tissue, with higher EZH2 levels significantly associated with higher grade gliomas (Zhang et al., 2015). To investigate whether the E2 and G7 cell lines also have increased expression of EZH2 or other PRC2 components, RNA-seq data from E2 and G7 cells was compared to data from nine normal neural stem cell lines (NSCs) grown under similar conditions (Mack et al 2019) (supplementary tables 3 and 4 and supplementary figure 1). Although expression of EZH2 itself was not significantly altered, the expression of two other members of the PRC2 complex, SUZ12 and RBBP7, was increased in both E2 and G7 cells. G7 cells also had increased expression of EZH1 (Figure 1b).

As both GSC cell lines have increased expression of PRC2 complex components, we hypothesised that they would be sensitive to inhibition of EZH2. The two lines have different cancer driver mutations, and represent different GBM subtypes, so comparing the effects of EZH2 inhibitors on both lines could provide valuable information about the future applications for EZH2 inhibitors. EZH2 inhibitors were chosen to test whether they can reduce the proliferation and survival of the GBM cell lines. Colony formation assays were used in the first instance, as they are considered to be the “gold standard” for assaying the efficacy of cytotoxic agents. Cells were plated at low density and treated with EPZ6438, GSK343 or UNC1999 for 48 hours. The compounds were then removed by media exchange and the cells were allowed to form colonies over a period of 14 days (Figure 2). All three compounds reduced colony formation by over 60% when used at concentrations of 2 μM or higher (p < 0.05). IC50 values were similar for all three inhibitors in both cell lines (1.27 – 1.9 μM), except for UNC1999 in E2 cells where the IC50 was 0.73 μM. These data show that even a relatively short exposure to an EZH2 inhibitor can have longer term effects on the ability of cells to survive and proliferate.

**Figure 2.**
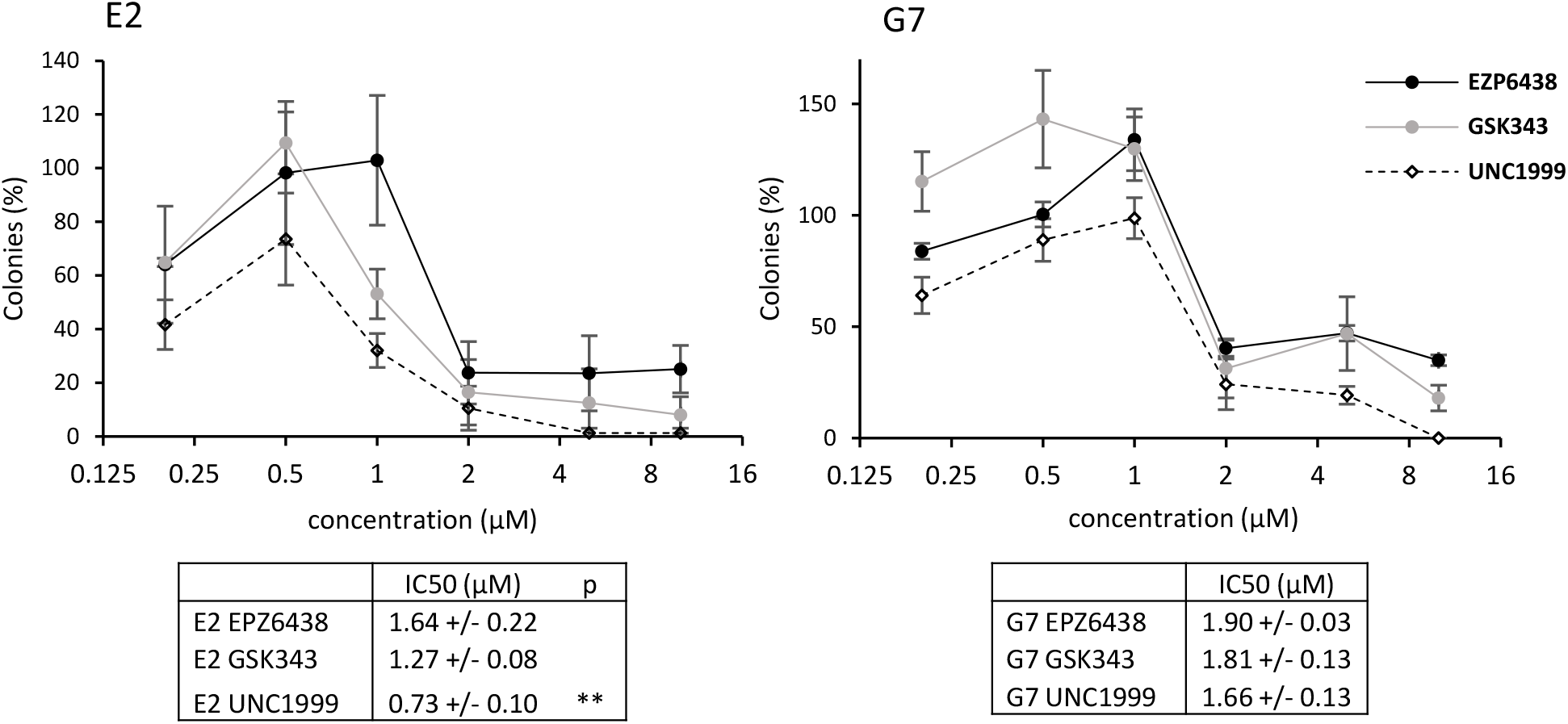
Treatment with EZH2 inhibitors reduces the clonogenic survival of GBM stem cells. Cells were plated at a low density, incubated with EPZ6438, GSK343, UNC1999 or DMSO for 48 hours, then allowed to form colonies over 14 days. Data points represent the mean and SEM of three independent experiments, as a percentage of DMSO treated control. The IC50 was calculated from each experiment, and the mean and standard deviation is shown. ** p < 0.01 for IC50 of E2 cells treated with UNC1999 compared to treatment with EPZ6438 or GSK343.

In order to investigate the effects of continuous exposure to EZH2 inhibitors on GBM cell line survival in the short term, a 96-well plate assay was used. After cells were cultured in the presence of EPZ6438, GSK343 or UNC1999 for five days, their relative cell density was determined (Figure 3). Both cell lines were sensitive to GSK343 and UNC1999, but were less sensitive to EPZ6438 (p < 0.05). Cells were consistently less sensitive to the EZH2 inhibitors in this assay than in the colony formation assay, as indicated by the higher IC50 values. For example, the IC50 values for GSK343 in E2 and G7 cells were 4.2 μM and 9.7 μM, respectively, in this short-term survival assay. The increased sensitivity to EZH2 inhibitors observed in the colony formation assays could reflect the requirement for treated cells to be able to proliferate several times to form a colony, or could be a result of the cells being more sensitive to the EZH2 inhibitors at low cell densities.

**Figure 3.**
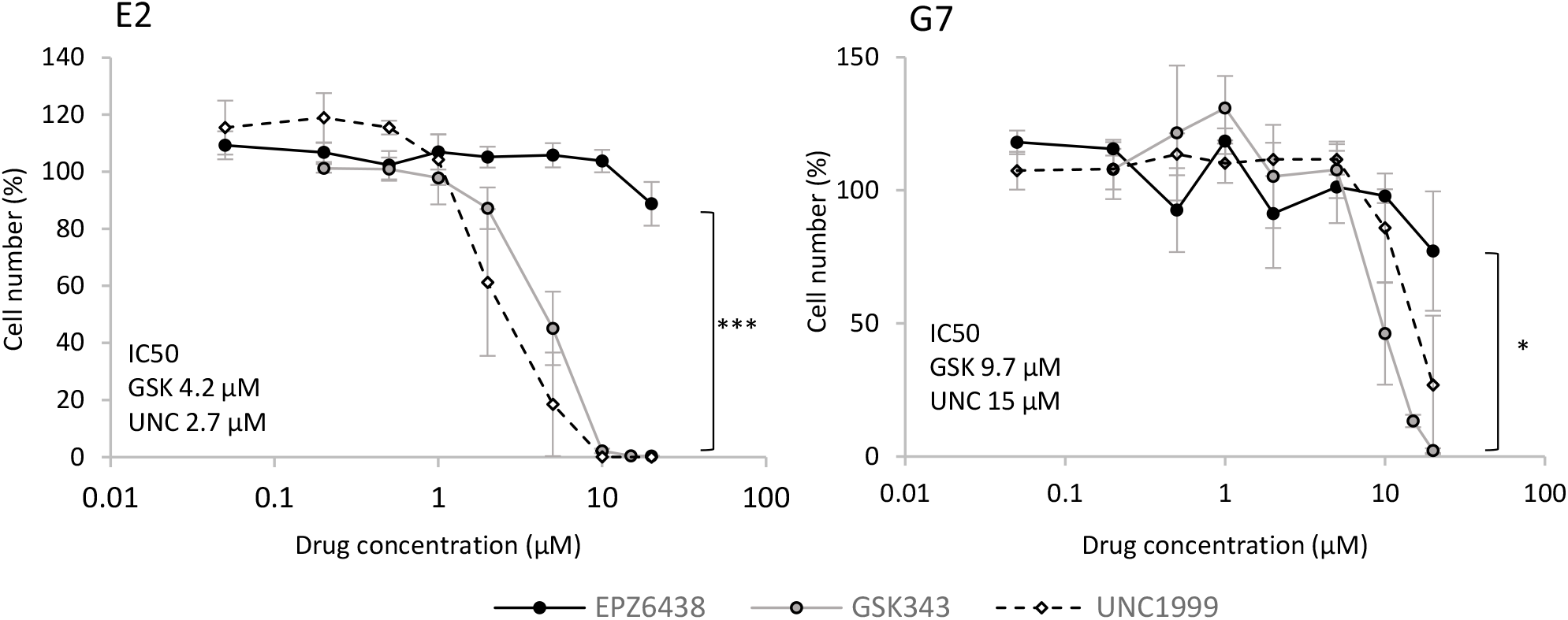
EZH2 inhibitors inhibit GBM stem cell proliferation. GBM cells were seeded in triplicate in 96 well plates in the presence of varying concentrations of EZH2 inhibitors and allowed to grow for five days before cell viability was assayed as a percentage of the DMSO control. Results are plotted as the mean value of three independent experiments. Error bars show the SEM. * p < 0.05. *** p < 0.001

### EZH2 inhibitors can sensitise GBM cells to radiation

Radiotherapy is a primary treatment for GBM. However, GSCs are more resistant to radiotherapy than the differentiated tumour cells that make up the bulk of the tumour (Carruthers et al., 2018). Therefore, there is considerable interest in identifying agents that can induce radio-sensitisation of these stem cells. Both GBM stem cell lines were treated with EZH2 inhibitors for 24 hours before and after irradiation, then allowed to form colonies. Irradiation of E2 cells with 5 Gy reduced the fraction of surviving colonies (surviving fraction, SF) to 0.36, whereas the combination of 5 Gy with 2 μM EZH2 inhibitor reduced the SF to 0.11 – 0.2 (Figure 4). Radiosensitisation can be quantified by calculating the dose modifying ratio (DMR), which is the ratio of the radiation dose required to reduce the SF to 0.37 in control versus treated samples (Carruthers et al., 2015). In E2 cells, the DMR for control vs EZH2 inhibitor ranged from 1.25 – 1.41 (p < 0.05), indicating that treatment with 2 μM inhibitor significantly increased the radiosensitivity of the cells. G7 cells were more sensitive to irradiation than E2 cells, with 5 Gy reducing the SF to 0.17 in control cells. Of the EZH2 inhibitors, only 2 μM GSK343 significantly radiosensitised the cells, with a DMR of 1.62. These data show that EZH2 inhibitors can induce radiosensitisation of GBM stem cells, but that the magnitude of this effect varies between cell lines. It is possible that radiosensitisation by EZH2 inhibitors is restricted to cells with low initial radiosensistivity.

**Figure 4.**
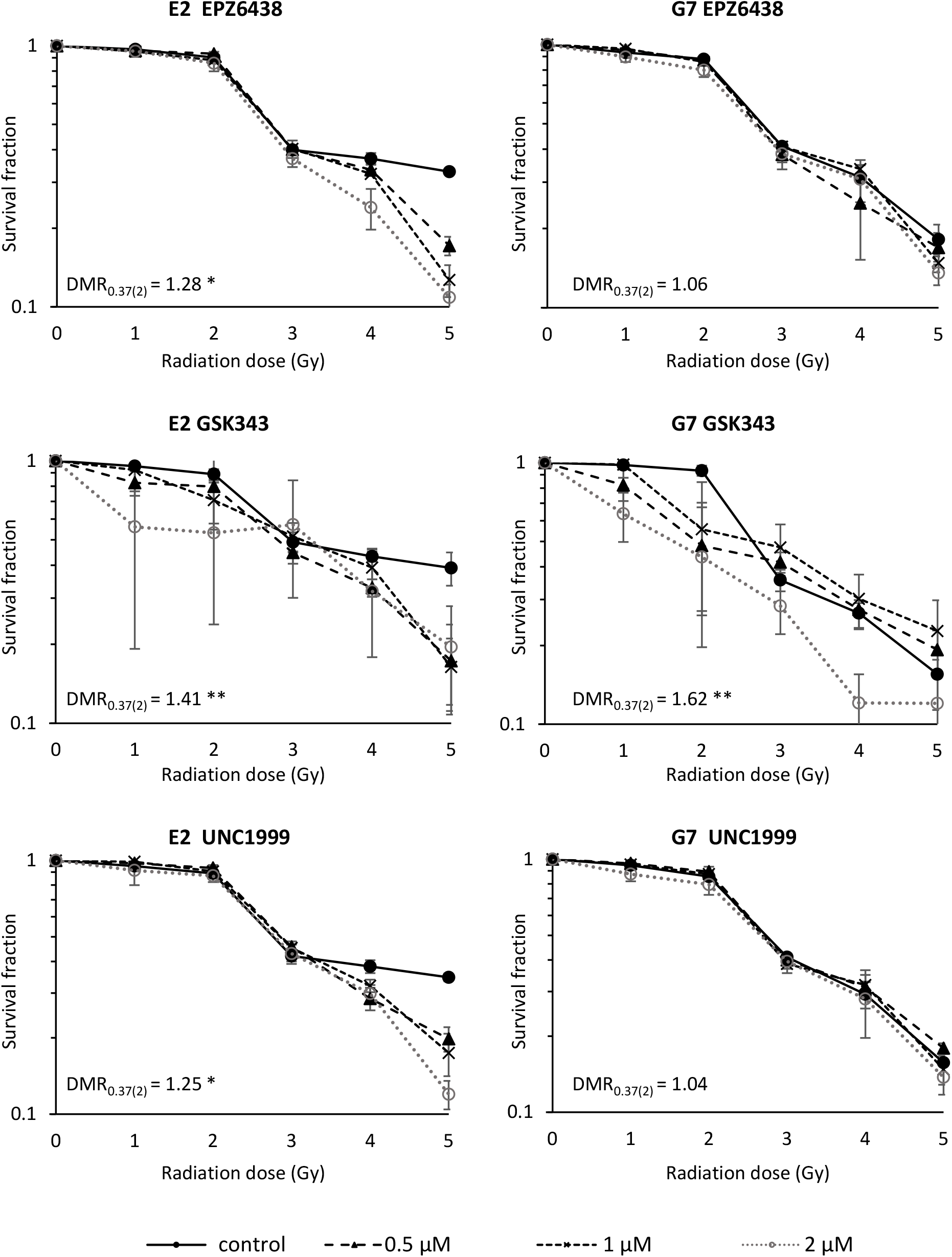
Clonogenic survival assay showing the combined effect of radiation with EZH2 inhibitors in GBM cells. Cells were incubated with different concentration of EZH2 inhibitor for 24hrs, exposed to radiation, then incubation in the presence of the drug was continued for a further 24hrs. The drug was then washed out and colonies allowed to form over 14 days. Results show the mean survival fraction (SF) from three independent colony assay experiments, error bars represent SEM. The dose modifying ratio (DMR) was calculated after regression modelling, and represents the dose required for SF 0.37 in control vs cells treated with 2 uM inhibitor. * p < 0.05, ** p < 0.01

### Inhibition of EZH2 leads to increased expression of neuronal genes

In order to investigate the molecular basis of how EZH2 inhibition reduces cell proliferation and survival, gene expression in treated cells was analysed using RNA-seq. Differentially expressed genes (DEGs) in drug treated cells were identified by pairwise comparison with the DMSO control (supplementary figure 2), with the Venn diagrams in Figure 5A showing the overlaps in DEGs between each EZH2 inhibitor. It is interesting to note that, although all three inhibitors induce similar reductions in colony formation and radiosensitisation, in E2 cells there are only 45 genes that are significantly changed in response to all three inhibitors. In contrast, in G7 cells there are only 572 DEGS common to all three inhibitors. Gene ontology was used to investigate which types of genes were altered, and the top five enriched GO categories in each experiment are shown in Figure 5b. E2 cells treated with UNC1999 only had 144 DEGs, which did not yield any significantly enriched GO categories. It can be seen that GO categories related to neurogenesis or neuronal structure are significantly enriched in all experiments shown.

**Figure 5.**
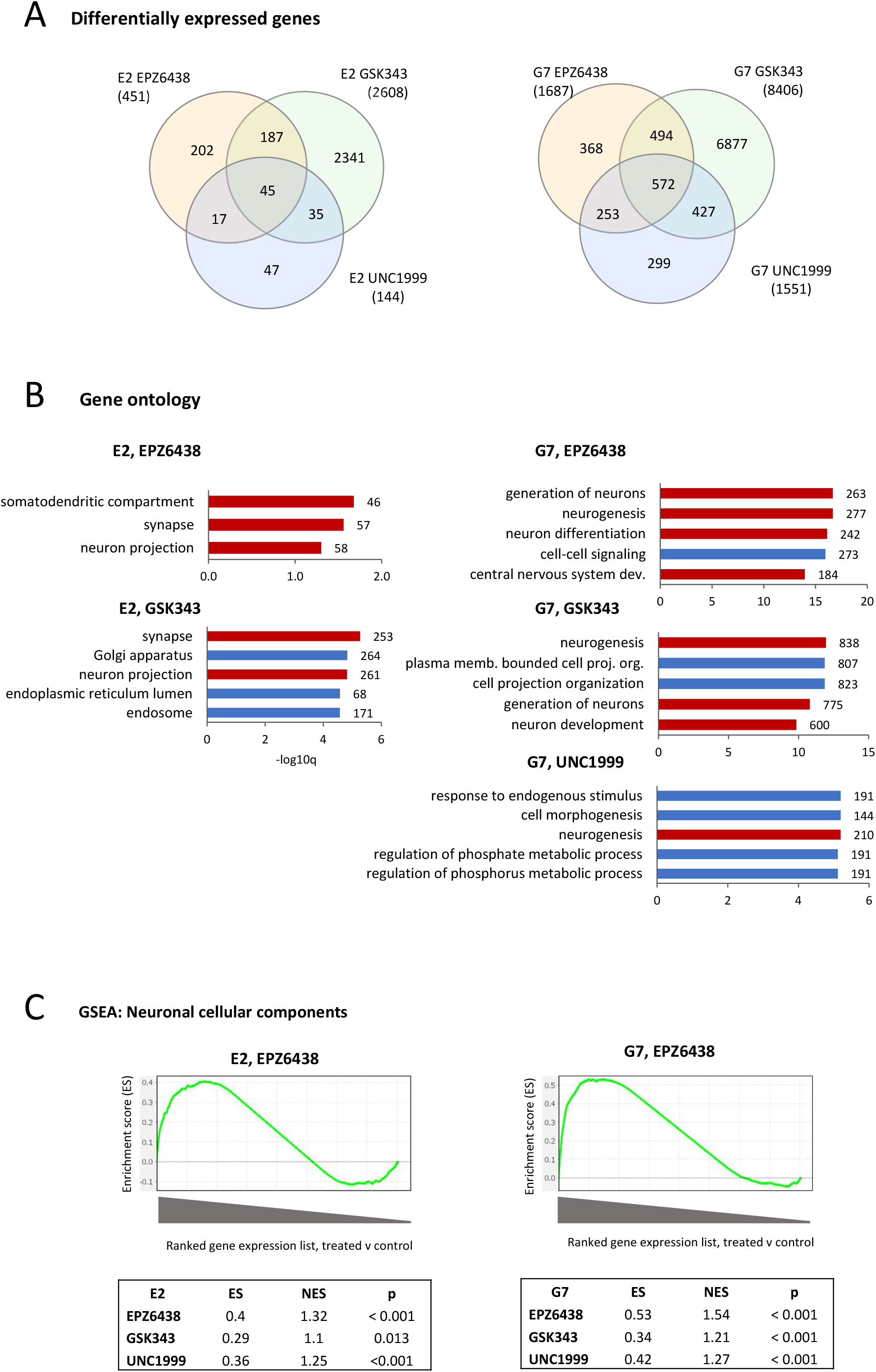
EZH2 inhibitors alter the expression of genes related to neurogenesis and neuronal structure. A) E2 and G7 cells were incubated with 2 μM EZH2 inhibitor for 5 days prior to RNA purification, RNA-seq and bioinformatic analysis. Differentially expressed genes in each experiment were identified by comparison to the DMSO control (adjp < 0.05), with the Venn diagrams showing the overlap in DEGs between each experiment (B) Gene ontology was performed on the list of DEGs from each experiment. The plots show the top five GO categories from each experiment. Red bars indicate GO categories with an explicit link to neuronal development or function. The x axis represents the -log10 of the Benjamini and Hochberg FDR value and bars are labelled with the number of genes from the DEG list. No GO categories were significantly enriched for E2 cells treated with UNC1999, probably due to the lower number of DEGs. Only three GO categories were enriched for E2 cells with EPZ6438. (C) GSEA analysis of changes in gene expression compared to a gene set of neuronal cellular components. Graphs show the running enrichment score across the ranked gene expression list, with genes most highly expressed in EPZ6438-treated cells compared to the control on the left. Tables show the enrichment score (ES), normalised enrichment score (NES) and p value for each experiment. The ES peak towards the left-hand side of each graph, and the significant p values, show that expression of neuronal cellular component genes is significantly higher in cells treated with EZH2 inhibitors than in the controls.

Gene set enrichment analysis (GSEA) is an alternative approach to identifying families of significantly enriched genes. It does not rely on a list of significant DEGs, but analyses the whole RNAseq dataset, so can detect more subtle changes across a whole family of genes. GSEA showed that expression levels of neuronal cellular components are significantly increased in both E2 and G7 cells treated with EPZ6438, GSK343 or UNC1999 (Figure 5c).

Taken together, the GO analyses and GSEA indicate that all three EZH2 inhibitors led to increased expression of neuronal genes in both GBM cell lines, which could be indicative of neuronal differentiation. In order to investigate whether there are common regulatory factors in all six experiments that might be driving these increases in neuronal gene expression, the RNAseq data were re-analysed by using group comparisons in which all the EZH2 inhibitor datasets were compared to the control. DEGs common to both E2 and G7 analyses were then identified (Figure 6a and b). This yielded a list of 147 common DEGs, 118 of which were upregulated in both E2 and G7 cells. Gene ontology revealed that these 118 genes are significantly enriched in categories related to neuronal development and neuronal structure (Figure 6c), and 48 of them have a known neuronal association (Figure 6d), consistent with the analysis of individual experiments.

**Figure 6.**
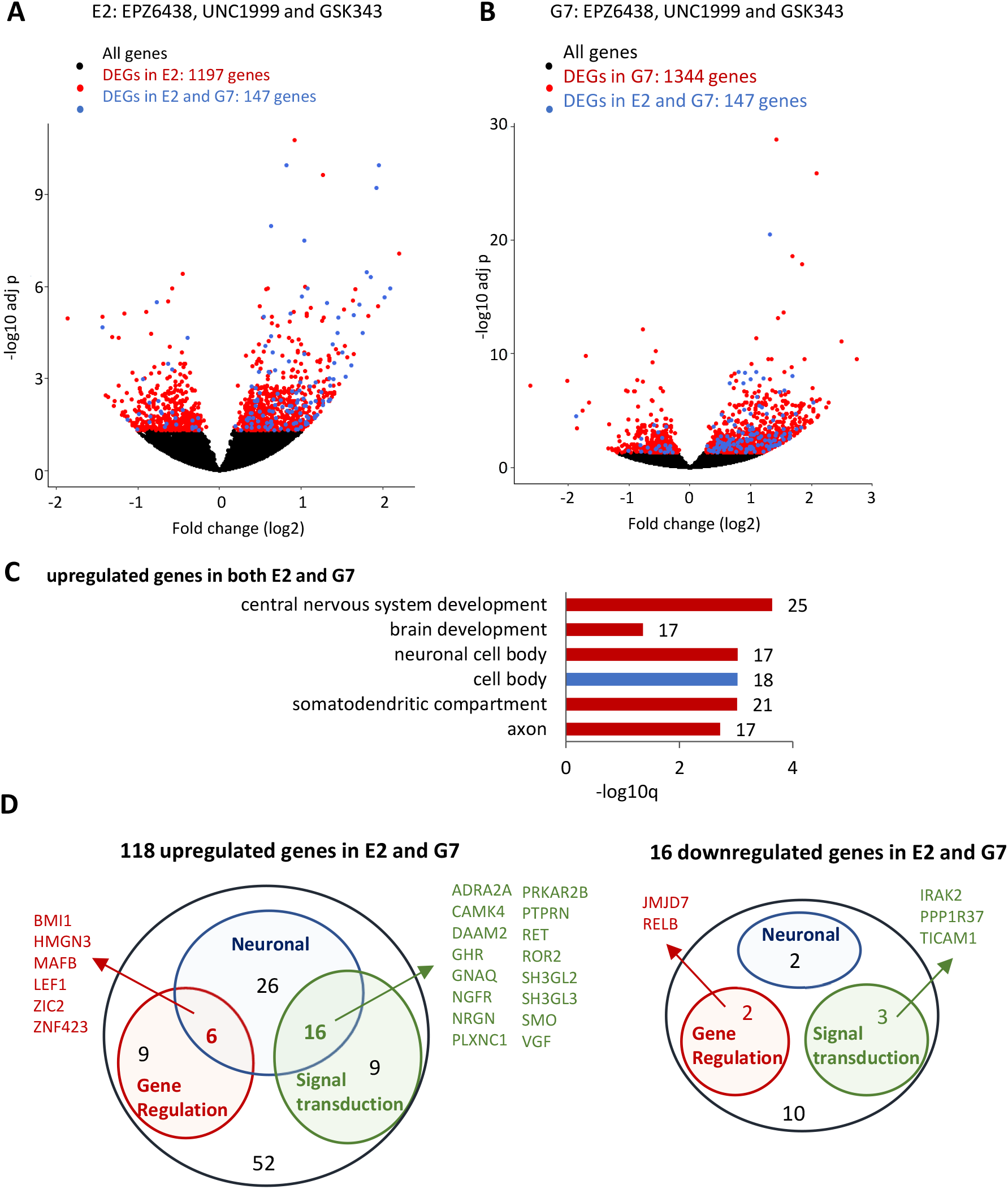
Combined RNA-seq analysis identifies a core list of DEGs common to both cell lines. A and B) Volcano plots of gene expression data from E2 and G7 cells. For each cell line, RNA-seq data from all samples treated with EZH2 inhibitors were compared to controls using DESEQ2, and DEGs (adjp < 0.05) were identified (red). The 147 DEGs common to both E2 and G7 cells are indicated in blue. 118 of these DEGs are upregulated in both E2 and G7 cells, 20 are downregulated in both lines, and 9 are differentially regulated. C) Upregulated DEGs common to both E2 and G7 cells were analysed for statistically enriched GO terms, and the top six GO terms are plotted as a bar chart. The x axis is -log10 of the B&H FDR, bars are labelled with number of DEGs in each category. D) Genes up or down regulated in both E2 and G7 cells are grouped according to whether they have a role in neuronal structure or function (indicated by GO terms), gene regulation and/or signal transduction.

To identify potential regulatory factors that might be driving the observed reduction in cell proliferation and the increase in neuronal gene expression, the list of DEGs was scanned for genes with roles in signal transduction or gene regulation (Figure 6d and additional table 1). Notably, the hedgehog pathway receptor component Smoothened (SMO), Wnt signalling regulator DAAM2 and the Wnt-activated transcription factor LEF1 are all upregulated. Hedgehog and Wnt signalling are well known to play many roles in regulating brain development and neurogenesis (De Luca et al., 2016, Bengoa-Vergniory and Kypta, 2015). Genes upregulated by EZH2 inhibitors could potentially be direct targets of EZH2’s H3K27 methylation activity. However, downregulated genes are more likely to be indirect targets, where EZH2 inhibition has led to upregulation of a repressor. One interesting example of a downregulated gene is the histone tail endopeptidase JMJD7, whose downregulated has previously been associated with reductions in proliferation and increased terminal differentiation (Liu et al., 2018b, Liu et al., 2018a, Liu et al., 2017).

**Table 1:** Direct and indirect EZH2 target genes that may regulate neuronal gene expression. To identify putative EZH2 direct target genes driving increased expression of neuronal genes, the following procedure was used. Starting with the 147 DEGs common to E2 and G7 that were identified earlier, this list was filtered for i) a decrease in normalised H3K27me3 signal of more than 1 in both E2 and G7 cells following treatment with EPZ6438 (19 genes) and ii) a known role in regulation of chromatin structure or gene expression, or a role in signal transduction (9 genes). Three indirect targets of interest are also shown. The data show the fold change in gene expression (log2) or ChIP enrichment in cells treated with EZH2 inhibitors versus the untreated control. #N/A: no peak identified. Genes are considered to be potential direct targets of EZH2 activity when their expression is increased and H3K27me3 enrichment decreased in both E2 and G7 cells after EZH2 inhibition. The full table of 147 genes is shown in additional table 1.

| gene | function | EZH2 target | RNA expression: drug treated vs control |  |  |  | ChIP seq data: EPZ6438 treated vs control |  |  |  |
| --- | --- | --- | --- | --- | --- | --- | --- | --- | --- | --- |
| | | | E2 log2fc | E2 adjp | G7 log2fc | G7 adjp | $\Delta$ H3K27me3 in E2 | $\Delta$ H3K27me3 in G7 | $\Delta$ H3K4me3 in E2 | $\Delta$ H3K4me3 in G7 |
| BMI1 | chromatin | direct? | 1.03 | 3.3E-08 | 1.09 | 4.4E-09 | -5.9 | -3.8 | 0.4 | 1.8 |
| DAAM2 | signalling | direct? | 1.06 | 0.031 | 1.43 | 0.001 | -6.4 | -3.9 | 4.3 | -0.7 |
| MAFB | transcription factor | direct? | 1.35 | 0.001 | 1.21 | 1.4E-08 | -55.7 | -9.8 | 9.2 | 14.3 |
| NGFR | signalling | direct? | 0.66 | 0.005 | 1.47 | 0.006 | -10.7 | -28.4 | 5.7 | 10.5 |
| PTPRN | signalling | direct? | 1.19 | 0.016 | 1.37 | 0.013 | -23.7 | -33.4 | 6.9 | 3.9 |
| RET | signalling | direct? | 0.84 | 0.049 | 1.44 | 0.011 | -12.7 | -70.2 | 9.5 | 3.3 |
| ZIC2 | transcription factor | direct? | 0.60 | 0.047 | 1.03 | 0.000 | -4.8 | -9.8 | 19.9 | 29.0 |
| PLXNC1 | signalling | direct? | 1.06 | 0.008 | 1.02 | 0.015 | -21.3 | -19.5 | 28.4 | 24.1 |
| ZNF423 | transcription factor | direct? | 1.09 | 0.020 | 1.65 | 0.002 | -49.6 | -53.4 | -4.5 | 10.0 |
| JMJD7 | chromatin | indirect | -1.44 | 2.1E-05 | -1.87 | 3.3E-05 | #N/A | #N/A | -11.9 | -1.5 |
| SMO | signalling | indirect | 0.39 | 0.002 | 0.46 | 0.008 | #N/A | #N/A | -9.2 | -5.9 |
| LEF1 | transcription factor | indirect | 0.63 | 1.1E-08 | 1.74 | 2.1E-05 | #N/A | -12.7 | -20.4 | 9.7 |

### Epigenetic profiling reveals genes that are putative targets of EZH2 activity

We reasoned that the genes that are directly affected by inhibition of EZH2’s histone methyltransferase activity would show a decrease in H3K27 methylation at their promoters and have a corresponding increase in mRNA expression. An increase in the active promoter mark H3K4me3 might also be detected.

Chromatin immunoprecipitation followed by high throughput sequencing (ChIP-seq) was used to study the genome-wide profiles of H3K27me3 and H3K4me3 in E2 and G7 cells before and after treatment with EPZ6438 (supplementary figure 3). Peaks of histone H3K27me3 modification in untreated cells were called using MACS2; the quantification of ChIP-seq reads at these peaks in EPZ6438-treated versus control cells is shown in Figure 7a. Peaks that overlap the promoter or transcription start site (TSS) of genes upregulated in EPZ6438-treated cells are shown in blue. It can be seen that stronger H3K27me3 peaks are less intense in EPZ6438-treated cells than in the control, consistent with inhibition of EZH2 activity across the genome. H3K27 enrichments are decreased in the peak regions across the whole genome dataset in both cell lines following drug treatment. This is especially pronounced in genes that had the most H3K27me3 signal prior to treatment. However, H3K4me3 enrichments were generally similar before and after treatment, with a few exceptions (Figure 7a).

**Figure 7.**
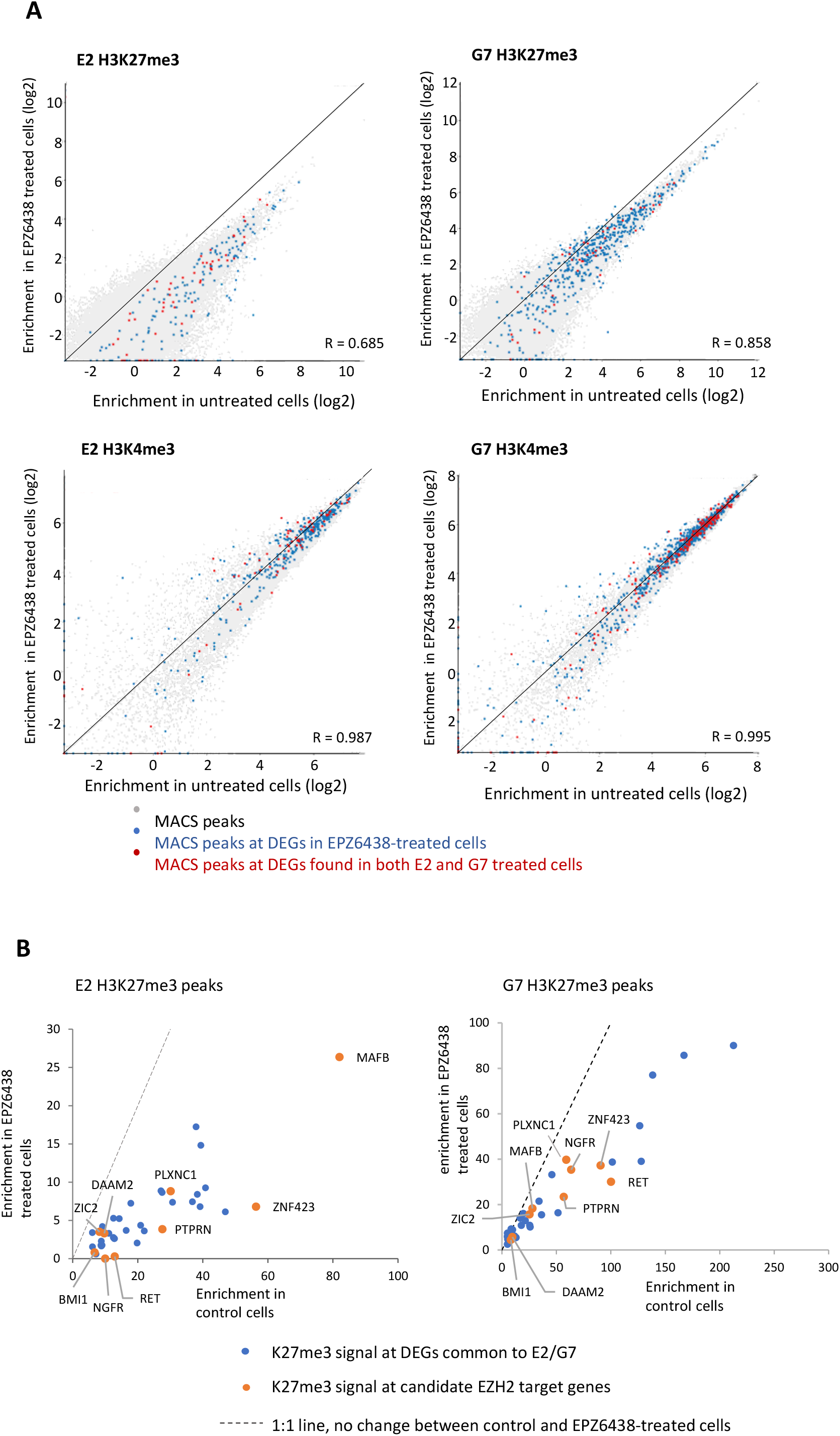
Identifying EZH2 target genes by epigenetic profiling. A) Scatter plots of H3K27me3 and H3K4me3 enrichment in control and EPZ6438-treated E2 and G7 cells. H3K27me3 enrichment was quantified at peaks that had been defined using MACS2 in control E2 and G7 cells. H3K4me3 enrichment was quantified at regions +/- 1500 bp from transcription start sites. Quantified regions that overlap with the promoter or transcription start sites (TSS) of DEGs that are upregulated in EPZ6438-treated cells are indicated in blue. Regions that overlap with the promoter or TSS of the 147 DEGs common to both E2 and G7 cells treated with all three inhibitors are indicated in red. 1:1 diagonal lines indicate the location of equal counts between treated and untreated. Data are plotted on a log2 scale. B) Scatter plot showing the quantification of H3K27me3 peaks mapping to DEGs common to E2 and G7 cells. Peaks within 1500 bp of the TSS were included; 38 peaks in E2 cells and 39 peaks in G7 cells meet this criterion. Nine candidate EZH2 target genes that show a decrease in K27me3 signal in both E2 and G7, and have functions related to gene expression or signal transduction are shown in orange and labelled (listed in table 1).

In order to identify which of the 147 DEGs common to both E2 and G7 might be direct targets of EZH2, we asked which of these genes had H3K27me3 peaks near the transcription start site (additional table 1). Most of these 147 genes did not have a H3K27me3 peak, so are not considered to be direct EZH2 targets. H3K27me3 peaks overlapping one of these DEGs are indicated in red in Figure 7a and are plotted separately in Figure 7b (38 and 39 peaks in E2 and G7 cells, respectively). Nearly all of these DEGs had reduced H3K27me3 signal and were upregulated following EZH2 inhibition (Figure 7b and additional table 1), indicating they may be direct targets of EZH2 activity.

To identify gene targets of EZH2 activity that are common to both GBM stem cell lines, DEGs that showed a reduction in H3K37me3 signal in both cell lines following EPZ6438 treatment were selected (additional table 1). Genes with roles in signal transduction, chromatin or gene regulation were extracted from this list in order to focus on regulatory factors that could driving broad changes in cellular phenotype (table 1). Figure 7b (orange points) shows that H3K27me3 signal is reduced at these nine genes following EZH2 inhibition. Interestingly, most of these nine genes also showed an increase in the active mark H3K4me3 in both cell lines (table 1).

The nine genes identified by this selection procedure comprise potential EZH2 targets whose expression is increased following EZH2 inhibition, and that may play roles in driving the observed increase in expression of neuronal genes in both cell lines. Genes of particular interest include the transcription factors MAFB, ZIC2 and ZNF423, all of which have key roles in regulating brain development and neuronal differentiation (Harder et al., 2014, Frank et al., 2015, Pai et al., 2020). Figure 8 shows the genomic regions for MAFB and ZIC2, and the region around the TSS of ZNF423. The EPZ6438-treated samples clearly show a reduction in H3K27me3 signal and narrower MACS2 peaks, consistent with the increased expression of these genes.

**Figure 8.**
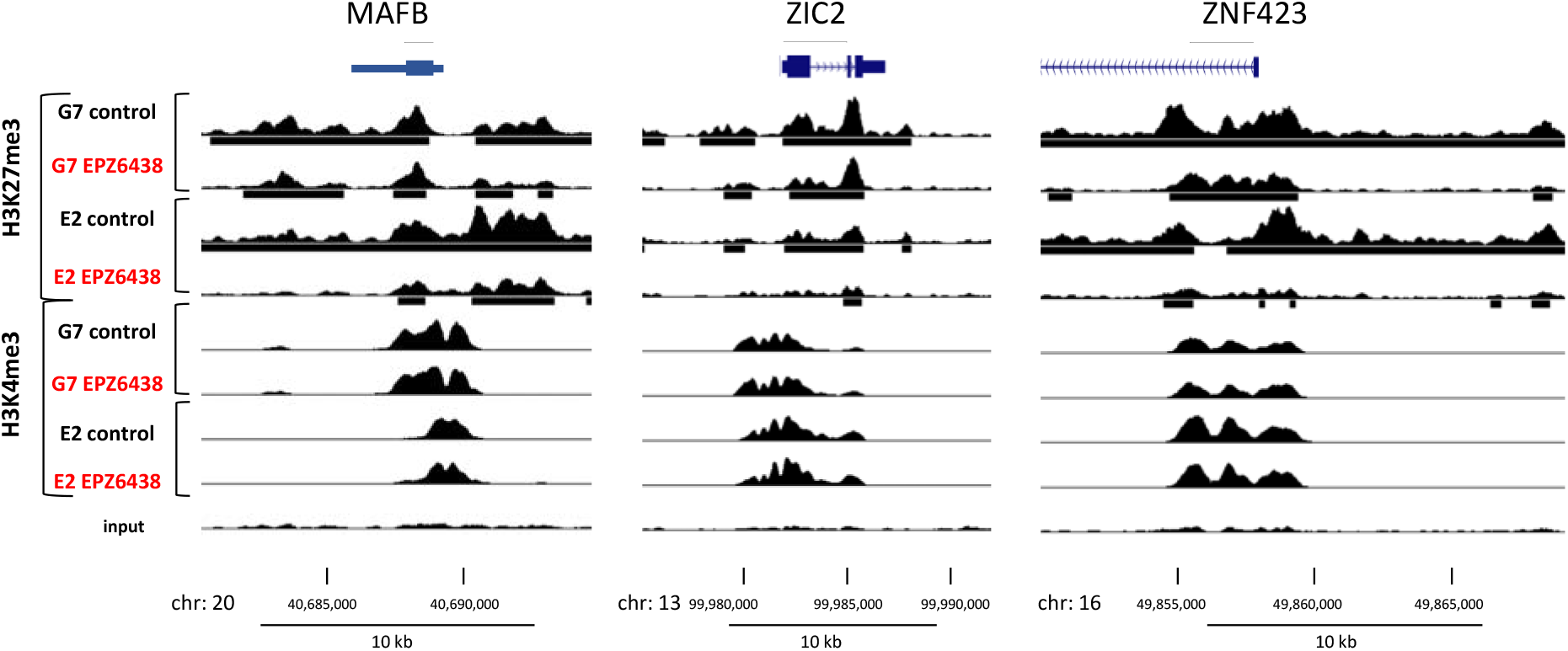
EPZ6438 induces epigenetic changes at key transcription factor loci. Visualisation of ChIP-seq signal and peaks at the MAFB, ZIC2 and ZNF423 gene loci. MACS2 was used to call peaks for each sample with comparison to the relevant input control using the broad peak setting. The input track shown is the E2 control and is representative of all input tracks generated. Y axis maxima for H3K27me3: 90, H3K4me3: 600. Input: 110

## Discussion

In this study, we have investigated the transcriptional and epigenomic outcomes of treating two GSC lines with three EZH2 inhibitors, with the aim of identifying common gene targets of EZH2’s histone methylation activity.

Using two GSC lines that correspond to different GBM subtypes has allowed us to focus on responses to EZH2 inhibitors that are common to both cell lines, and thus are more likely to translate to a wider range of GBM stem cells. Specifically, gene expression profiling revealed that E2 GSCs have similarities to both the classical and pro-neural GBM subtypes, whereas G7 cells are more mesenchymal (Verhaak et al., 2010, Brennan et al., 2013). It is important to note that although GBM subtypes have significant associations with some genetic changes (Brennan et al., 2013, Verhaak et al., 2010), transcriptional profiling has shown that several subtypes can be present in the same glioblastoma tumour, with proneural GBM stem cells associated with vascular regions and mesenchymal cells with hypoxia (Jin et al., 2017). Therefore, the gene expression profile of GBM stem cells is likely to represent a combination of their genetic, epigenetic and environmental influences.

Both GSC lines were sensitive to EZH2 inhibition, with both increasing their expression of the PRC2 complex components SUZ12 and RBBP7 in an effort to compensate. Although it might be expected that epigenetic inhibitors would require long term exposure in order to observe an effect (Knutson et al., 2014), it was notable that a 48 hr exposure to EZH2 inhibition is sufficient to observe a significant decrease in subsequent colony formation, with IC50 values in the region of 0.73 – 1.9 μM. The GSCs are less sensitive to the EZH2 inhibitors during a five day continuous exposure cell survival assay, however, implying that the sensitivity of GSCs to EZH2 inhibitors may be cell density-dependent. All three EZH2 inhibitors are able to sensitise E2 cells to radiation, even at the lowest concentration of 0.5 μM. However, it was recently demonstrated that a 3D culture system for GSCs was more accurate at predicting the clinical efficacy of radiosensitisation strategies than traditional 2D systems, so it will be important to assess the radiosensitisation following EZH2 inhibition under 3D conditions (Gomez-Roman et al., 2017, Gomez-Roman et al., 2020).

RNA-seq was performed to investigate whether there is a common molecular basis for the effect of EZH2 inhibition on GSC survival and proliferation. Strikingly, there was little overlap between the lists of differentially expressed genes following treatment with each inhibitor. However, gene ontology analysis and GSEA revealed that genes involved in neurogenesis and/or neuronal structure and function were enriched in all the experiments. Further analysis of the data revealed a subset of 147 common genes whose expression was changed in both cell lines following treatment with any of the three EZH2 inhibitors. These genes were also highly enriched in neuronal ontologies, indicating that increased neuronal gene expression is a common outcome of inhibiting EZH2 in GSCs.

We then asked whether there were any direct targets of EZH2 histone methylation activity that could be driving the increase in neuronal gene expression in these experiments. Integration of ChIP-seq data with the RNA-seq data allowed us to identify genes whose increased expression was accompanied by a decrease in H3K27me3 levels in both GSC lines. Nine of these have roles in regulating gene expression or signal transduction, including ZNF423, ZIC2, MAFB and BMI1.

ZNF423/ZFP423 is a 30-zinc finger transcription factor that is required for the development of the cerebellum, and for forebrain and hindbrain midline patterning (Alcaraz et al., 2006, Harder et al., 2014). Within the cerebellum, it is important for Purkinje cell differentiation and granule cell proliferation (Cheng et al., 2007). ZNF423 interacts with signalling molecules in several different pathways, including Notch, BMP, retinoic acid and SHH, thus integrating a range of signals to regulate transcription during both embryonic development and carcinogenesis (Harder et al., 2014, Hong and Hamilton, 2016).

Notably, ZNF423 has previously been shown to be a target of PRC2-dependent H3K27 tri-methylation in a mouse model of gliomagenesis (Signaroldi et al., 2016). Silencing of ZNF423 by H3K27me3 increased the activity of transcriptional networks relating to de-differentiation and cell migration, and was associated with decreased survival in both the mouse model and in human GBM patients (Signaroldi et al., 2016). Our data showing that pharmacological inhibition of EZH2 in human GBM stem cells leads to de-repression of ZFP423 and increased expression of neuronal genes is consistent with the work of Signaroldi and colleagues and reinforces the importance of the EZH2-ZNF423 axis in GBM development.

ZIC2 is a zinc finger transcription factor that is essential for brain development; homozygous ZIC2 mutations are frequently associated with holoprosencephaly, a severe forebrain malformation disorder (Barratt and Arkell, 2018). ZIC2 binds to enhancer elements in a developmentally regulated manner and has been shown to be required for the normal proliferation of cerebellar granular neuron precursors, and for appropriate gene expression in mature, postmitotic cerebellar granular neurons (Frank et al., 2015). In mouse embryonic stem cells, ZIC2 acts with the Mbd3-NURD chromatin remodelling complex at enhancers and is essential for ESC proliferation and neural differentiation (Luo et al., 2015). Interestingly, mathematical modelling of human brain gene expression using data from the Allen brain atlas has revealed that EZH2 is a key regulator of gene expression in the cerebellum, and that ZIC2 is one of the predicted EZH2 targets (Hecker et al., 2017), which is consistent with our findings presented here.

MAFB is a basic leucine zipper transcription factor that is essential for the correct development of the hindbrain (Sturgeon et al., 2011, Cordes and Barsh, 1994). It is also a neuronal lineage-specific regulator that is critical for the formation of specialised synapses in the auditory circuit of the inner ear (Yu et al., 2013), and participates in the regulation of cortical interneuron generation, maturation and synaptogenesis (Pai et al., 2019, Pai et al., 2020).

The most significantly upregulated gene across all the EZH2 inhibition experiments is the PRC1 complex component, BMI1, with a corresponding decrease in H3K27me3 levels at the BMI1 gene. The data indicate that EZH2 negatively regulates BMI1 expression, and we speculate that this may help to maintain the co-ordinated regulation of PRC-dependent gene silencing in GSCs. This contrasts with evidence from bronchial epithelial cells and hepatocellular carcinoma cells in which EZH2 has been shown to positively regulate production of BMI1 protein by repressing microRNA-218 and microRNA-200c expression, respectively (Wang et al., 2016, Xu et al., 2020). There is considerable evidence demonstrating a role for BMI1 in CNS development and NSC proliferation (Desai and Pethe, 2020), but further research would be required to investigate whether increased BMI1 can drive neuronal gene expression in the context of EZH2 inhibition.

Indirect, secondary targets will also be important in regulating the response to the EZH2 inhibition, along with proteins that are targets of EZH2’s non-histone dependent activity (Kim et al., 2013, Yi et al., 2021). For example, components of the Wnt and Sonic hedgehog (Shh) signalling pathways were highlighted by our analyses. Specifically, the Wnt signalling regulator DAAM2 is upregulated and is direct target of EZH2 activity in both GSC lines. LEF1, a transcription factor regulated by Wnt signalling, is upregulated in both lines, and appears to be a direct EZH2 target in G7 cells, but is not a target in E2 cells as no H3K27 methylation was detected. SMO, a component of the Shh receptor, is also upregulated following EZH2 inhibition, but is not a direct target of EZH2 H3K27 methyltransferse activity. Shh signalling plays key roles in the development of the central nervous system, particularly in the formation of the cerebellum (De Luca et al., 2016). Wnt signalling is also crucial for many aspects of brain development and neurogenesis, and its effects are highly context dependent and depend on the cell type and developmental stage being studied (Bengoa-Vergniory and Kypta, 2015).

Another indirect target of EZH2 activity, JMJD7, is worth noting. JMJD7 has been reported to possess endopeptidase activity that can cleave arginine methylated histone H3 and H4 N-terminal tails (Liu et al., 2017). It has been proposed that this activity of JMJD7 (and that of the homologous protein JMJD5) could promote transcription by creating tailless nucleosomes just after the transcriptional start sites, thus allowing paused RNA polymerase II to continue with elongation (Liu et al., 2017, Shen et al., 2017). A counter to this proposal perhaps, is the observation that JMJD7 does not engage with the H3 or H4 tails if they carry lysine acetylation of methylation, both of which are highly enriched at active TSS (Liu et al., 2018a).

It has been shown that JMJD7 is downregulated during osteoclast differentiation (Liu et al., 2018b), and that knockout of JMJD7 greatly decreases the ability of breast cancer cells to form colonies (Liu et al., 2017). In our study, JMJD7 expression is decreased following EZH2 inhibition, and, based on these previous studies, we can speculate that loss of JMJD7 activity may be an important contributing factor in the reduction of GSC colony formation. A recent study has identified a JMJD7 inhibitor, Cpd-3, that inhibits proliferation in several types of cancers, and it would be interesting to investigate whether Cpd-3 shows activity against GSC lines (Zhang et al., 2021)

In summary, we have identified a common set of genes whose expression is altered following EZH2 inhibition in pro-neural/classical and mesenchymal GSCs. It should be noted that additional research is needed see if these results can be replicated in other GSC lines, as genetic and epigenetic variability is known to influence the response to EZH2 inhibitors (Knutson et al., 2014, Signaroldi et al., 2016, McCabe et al., 2012). Our analyses revealed three neuronal transcription factors, ZNF423, ZIC2 and MAFB as direct targets of EZH2 activity, which are strong candidates for driving an increase in neuronal gene expression in GSCs following EZH2 inhibition. Loss of JMJD7 expression may also be a key factor in reducing GSC proliferation, opening up the potential for testing JMJD7 inhibitors in GSCs.

## Acknowledgements

We thank the Anthony Chalmers and members of his laboratory for training and advice in using the GSC lines, and we thank members of Adam and Katherine West’s groups for their advice and support throughout this project. We also thank the many undergraduate and postgraduate students who have assisted in the testing, development and replication of the methods and analyses used in this study. Adam West is funded by the BBSRC (BB/J008605/1) and MRC (MR/R005567/1).

## Material and Methods

### Cell culture and assays

E2 and G7 cells were a kind gift from Anthony Chalmers laboratory, originally derived from anonymised patient resection specimens as described previously (Fael Al-Mayhani et al., 2009, Carruthers et al., 2015). Cells were grown on Matrigel coated plates in AdDMEM F12 medium supplemented with 20ng/ml EGF, 10ng/ml bFGF,1% B27, 0.5% N2, 4ng/μl Heparin, and L-Glutamine (Fael Al-Mayhani et al., 2009, Gomez-Roman et al., 2017). EZH2 inhibitors EPZ6438, GSK343 and UNC1999 were from Selleckchem and dissolved in DMSO. Controls contained 0.2% DMSO. For colony assays, cells were plated at 250 or 500 cells/6 well plate (26 – 52 cells/cm^2^) exposed to EZH2 inhibitors or DMSO for 48 hours, then the inhibitors were removed and colonies allowed to grow for 14 days, prior to staining with crystal violet for counting (Carruthers et al., 2018). For irradiation colony assays, cells were plated in the presence of inhibitors for 24 hours, irradiated, then cultured in the same medium for another 24 hours. Irradiation was performed in a regularly calibrated x-ray cabinet (2.47 Gy/min, 195 kV, 15 mA, 0.5 mm Cu filter, X-Strahl)(Gomez-Roman et al., 2020). The media was removed and replaced with fresh media lacking inhibitors, and colonies were allowed to grow as described above. For proliferation assays, cells were plated at 2500 cells per well of a 96 well plate (7813 cells/cm^2^). After 24 hours, inhibitors were added, and plates were incubated for 5 days before being assayed using the Cell Titer-Glo kit (Promega).

Irradiation survival data were fitted using quadratic modelling; all modelled curves had R^2^ greater than 0.95 and F test p value of less than 0.01. The dose modifying ratio is the ratio of the irradiation dose required to reduce the surviving fraction to 0.37 in control vs 2 uM drug treatment (Carruthers et al., 2015). P values were calculated using 1-way ANOVA.

### RNA-seq

For RNA-seq, cells were plated in triplicate at 6 × 10^5^ cells per 10 cm dish and incubated with 2 μM inhibitors or DMSO for 5 days. RNA was purified using RNeasy kits (Qiagen); non-stranded library preparation and paired end sequencing (100 bp reads) of triplicate samples was performed by BGI Genomics (Hong Kong). After the quality of the data was checked using FASTQC within the Galaxy platform (Afgan et al., 2018), reads were aligned to Hg38 with HISAT2 (Kim et al., 2015) and transcripts were assembled using StringTie (Pertea et al., 2015), using the latest Hg38 GTF file (Ensembl) as a guide. DESeq2 (Love et al., 2014) was used to quantify transcripts. For comparison against gene expression in normal neural stem cells grown under similar conditions, RNAseq data from Mack and co-workers (Mack et al., 2019) was used (GEO accession SRP161553), and FASTQ files were processed using procedures for RF stranded libraries.

To identify sequence variants in coding regions, VCF files were generated using RNAseq data from control (untreated) cells and compared to the Hg38 reference sequence using FreeBayes. To identify mutations in cancer driver genes (Bailey et al., 2018), files were uploaded to Open-Cravat (Pagel et al., 2020), and filtered to only analyse missense variants, have a Chasm plus p value < 0.01 and Chasm plus GBM p value < 0.05 (Tokheim and Karchin, 2019). The variants were then filtered to remove those with a VCF PHRED score of less than 2, fewer than 15 total reads, or a variant allele frequency of less than 0.25.

For gene ontology using Toppgene (Chen et al., 2009), differentially expressed genes (DEGs) were defined as those with an adjusted p value less than 0.05, and a background gene list was created from all genes expressed in E2 and G7 cells. Intersects between DEG sets were calculated using the InteractiVenn webtool (Heberle et al., 2015). Gene set enrichment analysis (GSEA) (Subramanian et al., 2005) was performed, comparing normalised counts from individual replicates to a gene set of neuronal cellular components, which contained 2123 genes from the GO categories neuron projection (GO:0043005) and synapse (GO:0045202), downloaded from Ensembl BioMart (Howe et al., 2021).

To identify genes commonly regulated in all six experiments, DESEQ2 was performed comparing data from all inhibitor-treated samples with the DMSO control, for each cell line. The DEGs (adjp < 0.05) were compared between E2 and G7 cells, and the intersect resulted in a list of 147 DEGs common to both E2 and G7 cells. Additional table 1 shows the full list of 147 common DEGs, annotated with their functional category, expression data and Chip-seq data.

### ChIP-seq

Chromatin immunoprecipitation was performed as described previously (Garza-Manero et al., 2019) Anti-H3K4me3 and anti-H3K27me3 were obtained from EMD Millipore (07-473 and 07-449). Two to three ChIP samples were pooled prior to library preparation and paired end sequencing (150 bp reads) by Novogene. FASTQ files were aligned to Hg38 using Bowtie2 (Langmead and Salzberg, 2012).

To view the ChIP-seq data on the UCSC genome browser (Kent et al., 2002), the bamCoverage (Ramírez et al., 2016) tool was used to create BigWig files. The bin size was 50 bases and reads were normalised to reads to kilobase per million. BigWig files were uploaded to the Cyverse Discovery Environment, from where a public link was used to load the data into the UCSC genome browser. MACS2 was used to call peaks from ChIP samples using the data from input chromatin as the control (Zhang et al., 2008). The broad peak setting was used to combine nearby highly enriched regions into one broad region. Settings: band width 500 bases, q/FDR value of 0.05. The tool Chipseeker (Yu et al., 2015) was used to annotate the MACS output bed file with genomic regions derived from a Hg38 gtf file.

Seqmonk (Babraham Bioinformatics) was used to analyse the ChIP-seq peaks. H3K27me3 peaks were defined using the MACS2 peaks for the control (DMSO) sample in each experiment, and read counts across these probes were quantified in both the control and drug treated samples. For quantification purposes, H3K4me3 peaks were defined as regions spanning +/- 1500 bp from transcription start sites. Read counts were corrected per total million reads to allow for differences in library size and normalised to the input for that experiment. Peaks that overlapped the transcription start site (+/- 1500 bp) of upregulated differentially expressed genes were identified.

<u>Additional table 1 link</u>

