## Supplemental figures and tables for "EZH2 inhibition in glioblastoma stem cells increases the expression of neuronal genes and the neuronal developmental regulators ZIC2, ZNF423 and MAFB"

| GO Category | ID | Name | q-value FDR B&H | Hit Count in Query List | Hit Count in Genome |
|---|---|---|---|---|---|
| **enriched in E2** | | | | | |
| GO: Molecular Function | GO:0035254 | glutamate receptor binding | 5.74E-04 | 25 | 61 |
| GO: Molecular Function | GO:0005509 | calcium ion binding | 5.16E-03 | 139 | 685 |
| GO: Biological Process | GO:0022008 | neurogenesis | 4.44E-28 | 424 | 1817 |
| GO: Biological Process | GO:0048699 | generation of neurons | 9.29E-25 | 390 | 1687 |
| GO: Biological Process | GO:0030182 | neuron differentiation | 1.84E-22 | 355 | 1532 |
| GO: Biological Process | GO:0007267 | cell-cell signaling | 4.82E-19 | 391 | 1811 |
| GO: Biological Process | GO:0099536 | synaptic signaling | 8.92E-19 | 223 | 868 |
| GO: Biological Process | GO:0099537 | trans-synaptic signaling | 1.02E-18 | 219 | 849 |
| GO: Cellular Component | GO:0045202 | synapse | 1.33E-26 | 357 | 1476 |
| GO: Cellular Component | GO:0043005 | neuron projection | 2.93E-23 | 362 | 1567 |
| GO: Cellular Component | GO:0098794 | postsynapse | 3.22E-20 | 201 | 741 |
| GO: Cellular Component | GO:0036477 | somatodendritic compartment | 1.03E-17 | 254 | 1064 |
| GO: Cellular Component | GO:0097447 | dendritic tree | 5.05E-17 | 198 | 774 |
| GO: Cellular Component | GO:0030425 | dendrite | 5.05E-17 | 198 | 774 |
| **enriched in G7** | | | | | |
| GO: Molecular Function | GO:0005201 | extracellular matrix structural constituent | 1.36E-03 | 50 | 167 |
| GO: Molecular Function | GO:0005102 | signaling receptor binding | 3.06E-03 | 292 | 1533 |
| GO: Molecular Function | GO:0016773 | phosphotransferase activity, alcohol group as accep | 5.58E-03 | 167 | 814 |
| GO: Molecular Function | GO:0004672 | protein kinase activity | 8.32E-03 | 146 | 704 |
| GO: Molecular Function | GO:0016301 | kinase activity | 8.32E-03 | 175 | 873 |
| GO: Biological Process | GO:0045229 | external encapsulating structure organization | 1.16E-06 | 105 | 395 |
| GO: Biological Process | GO:0001501 | skeletal system development | 1.16E-06 | 133 | 539 |
| GO: Biological Process | GO:0030198 | extracellular matrix organization | 1.16E-06 | 104 | 393 |
| GO: Biological Process | GO:0043062 | extracellular structure organization | 1.16E-06 | 104 | 394 |
| GO: Biological Process | GO:0007155 | cell adhesion | 1.41E-06 | 288 | 1420 |
| GO: Biological Process | GO:0022610 | biological adhesion | 1.86E-06 | 288 | 1426 |
| GO: Biological Process | GO:0060429 | epithelium development | 1.11E-05 | 269 | 1340 |
| GO: Biological Process | GO:0061138 | morphogenesis of a branching epithelium | 3.06E-05 | 63 | 216 |
| GO: Biological Process | GO:0042127 | regulation of cell population proliferation | 5.29E-05 | 333 | 1754 |
| GO: Cellular Component | GO:0030312 | external encapsulating structure | 4.15E-08 | 141 | 565 |
| GO: Cellular Component | GO:0031012 | extracellular matrix | 4.15E-08 | 140 | 563 |
| GO: Cellular Component | GO:0062023 | collagen-containing extracellular matrix | 2.87E-05 | 107 | 444 |
| GO: Cellular Component | GO:0001650 | fibrillar center | 2.09E-03 | 39 | 131 |

**Supplementary table 1. Gene ontology analysis of E2 compared to G7 GBM.**

DESEQ2 was used to compare RNAseq data from E2 and G7 cells. Differentially expressed genes were selected with an adjusted p value of less than 0.01 and a log 2 fold change of greater than 1. The top six significantly enriched GO categories are shown, with a maximum FDR cut-off of 0.01. The number of genes in the RNAseq query set is shown, as is the total number of genes in the GO category.

| NAME | SIZE | ES | NES | NOM p-val | FDR q-val | RANK AT MAX | LEADING EDGE |
|---|---|---|---|---|---|---|---|
| **Enriched in E2 cells** | | | | | | | |
| HALLMARK_HEDGEHOG_SIGNALING | 36 | 0.60 | 1.50 | 0.01575 | 0.02449 | 4825 | tags=56%, list=12%, signal=63% |
| **Enriched in G7 cells** | | | | | | | |
| HALLMARK_TNFA_SIGNALING_VIA_NFKB | 200 | -0.59 | -1.81 | 0.00000 | 0.00073 | 5786 | tags=51%, list=14%, signal=59% |
| HALLMARK_MYC_TARGETS_V2 | 58 | -0.70 | -1.82 | 0.00000 | 0.00147 | 7178 | tags=67%, list=18%, signal=82% |
| HALLMARK_INFLAMMATORY_RESPONSE | 200 | -0.57 | -1.72 | 0.00000 | 0.00203 | 5786 | tags=43%, list=14%, signal=50% |
| HALLMARK_EPITHELIAL_MESENCHYMAL_TRANSITION | 200 | -0.56 | -1.70 | 0.00000 | 0.00236 | 3428 | tags=42%, list=9%, signal=45% |
| HALLMARK_MTORC1_SIGNALING | 200 | -0.58 | -1.73 | 0.00000 | 0.00270 | 5773 | tags=46%, list=14%, signal=53% |
| HALLMARK_CHOLESTEROL_HOMEOSTASIS | 74 | -0.59 | -1.58 | 0.00316 | 0.00697 | 6718 | tags=58%, list=17%, signal=70% |
| HALLMARK_UNFOLDED_PROTEIN_RESPONSE | 113 | -0.56 | -1.58 | 0.00000 | 0.00792 | 7405 | tags=53%, list=18%, signal=65% |
| HALLMARK_UV_RESPONSE_DN | 144 | -0.53 | -1.55 | 0.00000 | 0.01100 | 5476 | tags=44%, list=14%, signal=50% |
| HALLMARK_ALLOGRAFT_REJECTION | 200 | -0.49 | -1.47 | 0.00000 | 0.02416 | 6537 | tags=33%, list=16%, signal=39% |
| HALLMARK_HYPOXIA | 200 | -0.47 | -1.42 | 0.00142 | 0.03812 | 5153 | tags=39%, list=13%, signal=44% |
| HALLMARK_IL6_JAK_STAT3_SIGNALING | 87 | -0.51 | -1.40 | 0.01587 | 0.04043 | 6409 | tags=41%, list=16%, signal=49% |
| HALLMARK_APOPTOSIS | 161 | -0.47 | -1.40 | 0.00427 | 0.04228 | 6105 | tags=39%, list=15%, signal=46% |
| HALLMARK_ANDROGEN_RESPONSE | 100 | -0.50 | -1.38 | 0.02344 | 0.04359 | 4614 | tags=38%, list=11%, signal=43% |
| HALLMARK_INTERFERON_GAMMA_RESPONSE | 200 | -0.46 | -1.39 | 0.00289 | 0.04364 | 4008 | tags=29%, list=10%, signal=33% |
| HALLMARK_P53_PATHWAY | 200 | -0.45 | -1.37 | 0.00849 | 0.04503 | 7020 | tags=44%, list=17%, signal=53% |
| HALLMARK_XENOBIOTIC_METABOLISM | 200 | -0.45 | -1.37 | 0.00552 | 0.04560 | 4236 | tags=28%, list=11%, signal=31% |
| HALLMARK_IL2_STAT5_SIGNALING | 199 | -0.47 | -1.40 | 0.00429 | 0.04588 | 4285 | tags=31%, list=11%, signal=35% |

**Supplementary table 2. GSEA of gene expression in E2 cells versus G7 cells.**

Gene set enrichment analysis of changes in gene expression in E2 vs G7 cells, compared to the Hallmarks collection of genesets. Genesets with FDR q value < 0.05 are shown. Table columns: size: total number of genes in each gene set. ES: the enrichment score, NES: normalised enrichment score, which is the ES normalised for the size of the geneset, NOM p-val: nominal p value, FDR q-val: false discovery rate, controlled for the total number of genesets analysed. Rank at max gives the position in the ranked list at which the maximum ES was obtained. The leading edge refers to the subset of genes in the ranked list before the maximum ES is reached. "Tags" refers to the percentage of gene hits that are in the leading edge. "List" is the percentage of genes in the dataset that are in the leading edge. Signal is a combination of the previous two numbers, where a higher number indicates that the genes in the geneset are clustered closer towards the top of the ranked dataset.

| Category | ID | Name | FDR B&H | In query | In Genome |
|---|---|---|---|---|---|
| **E2 v NSCs upregulated DEGs** | | | | | |
| GO: Molecular Function | GO:0140110 | transcription regulator activity | 0.00001656 | 174 | 1821 |
| GO: Molecular Function | GO:0043565 | sequence-specific DNA binding | 0.0001812 | 155 | 1650 |
| GO: Molecular Function | GO:0000977 | RNA polymerase II transcription regulatory region sequence-spec | 0.0002082 | 136 | 1416 |
| GO: Molecular Function | GO:0001067 | transcription regulatory region nucleic acid binding | 0.0002082 | 144 | 1527 |
| GO: Molecular Function | GO:0000976 | transcription cis-regulatory region binding | 0.0002355 | 143 | 1523 |
| GO: Molecular Function | GO:0003700 | DNA-binding transcription factor activity | 0.0004005 | 129 | 1358 |
| GO: Biological Process | GO:0048706 | embryonic skeletal system development | 0.00003219 | 29 | 137 |
| GO: Biological Process | GO:0051254 | positive regulation of RNA metabolic process | 0.00005095 | 162 | 1707 |
| GO: Biological Process | GO:0048704 | embryonic skeletal system morphogenesis | 0.00008508 | 23 | 102 |
| GO: Biological Process | GO:0009792 | embryo development ending in birth or egg hatching | 0.00008508 | 88 | 790 |
| GO: Biological Process | GO:0009952 | anterior/posterior pattern specification | 0.00008508 | 36 | 217 |
| GO: Biological Process | GO:0008380 | RNA splicing | 0.0001135 | 57 | 440 |
| GO: Cellular Component | GO:0000785 | chromatin | 4.199E-09 | 129 | 1132 |
| GO: Cellular Component | GO:0005694 | chromosome | 1.302E-07 | 164 | 1649 |
| GO: Cellular Component | GO:0140513 | nuclear protein-containing complex | 0.0002374 | 115 | 1189 |
| GO: Cellular Component | GO:0005667 | transcription regulator complex | 0.0009265 | 56 | 483 |
| Domain | IPR020479 | Homeobox_metazoa | 4.637E-14 | 32 | 84 |
| Domain | PF00046 | Homeobox | 1.551E-12 | 48 | 203 |
| **E2 v NSCs downregulated DEGs** | | | | | |
| GO: Biological Process | GO:0006396 | RNA processing | 0.000008582 | 201 | 1242 |
| GO: Biological Process | GO:0006334 | nucleosome assembly | 0.00001904 | 32 | 98 |
| GO: Biological Process | GO:0031497 | chromatin assembly | 0.0001106 | 40 | 150 |
| GO: Biological Process | GO:0097549 | chromatin organization involved in negative regulation of transcr | 0.0003694 | 31 | 107 |
| GO: Biological Process | GO:0006333 | chromatin assembly or disassembly | 0.0006281 | 42 | 174 |
| GO: Biological Process | GO:0000353 | formation of quadruple SL/U4/U5/U6 snRNP | 0.0007133 | 8 | 10 |
| GO: Biological Process | GO:0000365 | mRNA trans splicing, via spliceosome | 0.0007133 | 8 | 10 |
| GO: Cellular Component | GO:0000786 | nucleosome | 1.869E-13 | 33 | 62 |
| GO: Cellular Component | GO:0044815 | DNA packaging complex | 9.572E-12 | 33 | 70 |
| GO: Cellular Component | GO:0005730 | nucleolus | 4.609E-11 | 222 | 1278 |
| GO: Cellular Component | GO:0032993 | protein-DNA complex | 0.00001915 | 44 | 176 |
| Domain | IPR016024 | ARM-type_fold | 3.597E-09 | 77 | 323 |
| Domain | IPR007125 | Histone_H2A/H2B/H3 | 1.249E-08 | 21 | 38 |
| **G7 v NSCs upregulated DEGs** | | | | | |
| GO: Molecular Function | GO:0008134 | transcription factor binding | 0.0002008 | 104 | 693 |
| GO: Molecular Function | GO:0140110 | transcription regulator activity | 0.0002924 | 224 | 1821 |
| GO: Biological Process | GO:0051254 | positive regulation of RNA metabolic process | 1.23E-05 | 220 | 1707 |
| GO: Biological Process | GO:0045944 | positive regulation of transcription by RNA polymerase II | 4.75E-05 | 166 | 1237 |
| GO: Biological Process | GO:0010557 | positive regulation of macromolecule biosynthetic process | 4.75E-05 | 229 | 1844 |
| GO: Biological Process | GO:1903508 | positive regulation of nucleic acid-templated transcription | 4.75E-05 | 204 | 1615 |
| GO: Biological Process | GO:0045893 | positive regulation of transcription, DNA-templated | 4.75E-05 | 204 | 1615 |
| GO: Biological Process | GO:1902680 | positive regulation of RNA biosynthetic process | 4.75E-05 | 204 | 1616 |
| GO: Cellular Component | GO:0000785 | chromatin | 1.91E-05 | 155 | 1132 |
| GO: Cellular Component | GO:0005694 | chromosome | 2.38E-04 | 203 | 1649 |
| GO: Cellular Component | GO:0035097 | histone methyltransferase complex | 2.80E-04 | 20 | 67 |
| Domain | IPR020479 | Homeobox_metazoa | 1.93E-08 | 31 | 84 |
| Domain | PF00046 | Homeobox | 2.03E-05 | 45 | 203 |
| **G7 v NSCs downregulated DEGs** | | | | | |
| GO: Molecular Function | GO:0030627 | pre-mRNA 5'-splice site binding | 3.46E-04 | 11 | 16 |
| GO: Biological Process | GO:0099111 | microtubule-based transport | 1.15E-08 | 123 | 532 |
| GO: Biological Process | GO:0007010 | cytoskeleton organization | 3.17E-07 | 283 | 1618 |
| GO: Biological Process | GO:0010970 | transport along microtubule | 3.17E-07 | 115 | 518 |
| GO: Biological Process | GO:0007018 | microtubule-based movement | 6.80E-07 | 136 | 656 |
| GO: Biological Process | GO:0098840 | protein transport along microtubule | 6.80E-07 | 98 | 428 |
| GO: Biological Process | GO:0099118 | microtubule-based protein transport | 6.80E-07 | 98 | 428 |
| GO: Cellular Component | GO:0000786 | nucleosome | 1.148E-13 | 35 | 62 |
| GO: Cellular Component | GO:0044815 | DNA packaging complex | 1.127E-12 | 36 | 70 |
| GO: Cellular Component | GO:0099081 | supramolecular polymer | 0.00007612 | 230 | 1368 |
| GO: Cellular Component | GO:0099080 | supramolecular complex | 0.00008893 | 270 | 1660 |
| GO: Cellular Component | GO:0099512 | supramolecular fiber | 0.0001561 | 226 | 1361 |
| GO: Cellular Component | GO:0032993 | protein-DNA complex | 0.0001751 | 45 | 176 |
| Domain | IPR007125 | Histone_H2A/H2B/H3 | 2.597E-10 | 24 | 38 |

**Supplementary table 3. Gene ontology analysis of GBM cells compared to normal neural stem cells.**
DESEQ2 was used to compare RNAseq data from E2 or G7 cells with a set of 9 normal neural stem cells. Differentially expressed genes were selected with an adjusted p value of less than 0.01 and a log 2 fold change of greater than 1 or less than -1. Up and down regulated DEGs were analysed separately using Toppgene. The top six significantly enriched GO categories are shown, with a maximum FDR cut-off of 0.001. The top significantly enriched protein domains are also shown. The number of genes in the RNAseq query set is shown, as is the total number of genes in the GO category.

| E2 vs NSCs, FDR < 0.05 | SIZE | ES | NES | NOM p-val | FDR q-val | RANK AT MAX | LEADING EDGE |
|---|---|---|---|---|---|---|---|
| HALLMARK_UV_RESPONSE_UP | 158 | 0.486023 | 1.764502 | 0 | 0.0028562 | 7961 | tags=46%, list=20%, signal=57% |
| HALLMARK_PANCREAS_BETA_CELLS | 40 | 0.615054 | 1.786074 | 0.00201207 | 0.004402 | 5068 | tags=28%, list=13%, signal=31% |
| HALLMARK_MYC_TARGETS_V1 | 200 | 0.402534 | 1.50539 | 0 | 0.0472493 | 8715 | tags=46%, list=22%, signal=58% |

| G7 vs NSCs, FDR < 0.05 | SIZE | ES | NES | NOM p-val | FDR q-val | RANK AT MAX | LEADING EDGE |
|---|---|---|---|---|---|---|---|
| HALLMARK_MYC_TARGETS_V2 | 58 | 0.750808 | 2.228056 | 0 | 0 | 6515 | tags=66%, list=16%, signal=78% |
| HALLMARK_TNFA_SIGNALING_VIA_NFKB | 200 | 0.552713 | 1.938636 | 0 | 0 | 5657 | tags=38%, list=14%, signal=43% |
| HALLMARK_MTORC1_SIGNALING | 200 | 0.547471 | 1.914119 | 0 | 0 | 6263 | tags=44%, list=16%, signal=51% |
| HALLMARK_UV_RESPONSE_UP | 158 | 0.527275 | 1.796855 | 0 | 0.0002949 | 5316 | tags=38%, list=13%, signal=44% |
| HALLMARK_INFLAMMATORY_RESPONSE | 200 | 0.510829 | 1.796816 | 0 | 0.0002359 | 6541 | tags=33%, list=16%, signal=39% |
| HALLMARK_UNFOLDED_PROTEIN_RESPONSI | 113 | 0.540108 | 1.776267 | 0 | 0.0007918 | 5977 | tags=42%, list=15%, signal=50% |
| HALLMARK_MYC_TARGETS_V1 | 200 | 0.499892 | 1.773503 | 0 | 0.0006786 | 6706 | tags=44%, list=17%, signal=53% |
| HALLMARK_CHOLESTEROL_HOMEOSTASIS | 74 | 0.562768 | 1.735682 | 0 | 0.0014071 | 3065 | tags=28%, list=8%, signal=31% |
| HALLMARK_ESTROGEN_RESPONSE_EARLY | 200 | 0.486211 | 1.715835 | 0 | 0.00193 | 6347 | tags=37%, list=16%, signal=44% |
| HALLMARK_ESTROGEN_RESPONSE_LATE | 200 | 0.477442 | 1.680411 | 0 | 0.002802 | 6347 | tags=32%, list=16%, signal=37% |
| HALLMARK_PANCREAS_BETA_CELLS | 40 | 0.599389 | 1.662198 | 0.00166389 | 0.0034317 | 2883 | tags=23%, list=7%, signal=24% |
| HALLMARK_ALLOGRAFT_REJECTION | 200 | 0.455324 | 1.598828 | 0 | 0.0073891 | 6166 | tags=22%, list=15%, signal=26% |
| HALLMARK_GLYCOLYSIS | 200 | 0.443877 | 1.553946 | 0 | 0.0111013 | 6435 | tags=34%, list=16%, signal=40% |
| HALLMARK_P53_PATHWAY | 200 | 0.43159 | 1.510718 | 0.00140056 | 0.0172107 | 6673 | tags=39%, list=17%, signal=47% |
| HALLMARK_REACTIVE_OXYGEN_SPECIES_PA | 49 | 0.516098 | 1.49987 | 0.01666667 | 0.018002 | 7916 | tags=47%, list=20%, signal=58% |
| HALLMARK_ADIPOGENESIS | 200 | 0.409728 | 1.446533 | 0.00542005 | 0.0322819 | 5831 | tags=31%, list=15%, signal=36% |
| HALLMARK_IL6_JAK_STAT3_SIGNALING | 87 | 0.444936 | 1.430051 | 0.01735016 | 0.0360786 | 5616 | tags=24%, list=14%, signal=28% |
| HALLMARK_IL2_STAT5_SIGNALING | 199 | 0.401188 | 1.415795 | 0.00408163 | 0.0401629 | 5868 | tags=25%, list=15%, signal=29% |

**Supplementary table 4. GSEA of gene expression in GBM cells versus normal NSCs.**

Gene set enrichment analysis of changes in gene expression in E2 or G7 cells versus normal NSCs, compared to the Hallmarks collection of genesets. Genesets with FDR q value < 0.05 are shown. Table columns: size: total number of genes in each gene set. ES: the enrichment score, NES: normalised enrichment score, which is the ES normalised for the size of the geneset, NOM p-val: nominal p value, FDR q-val: false discovery rate, controlled for the total number of genesets analysed. Rank at max gives the position in the ranked list at which the maximum ES was obtained. The leading edge refers to the subset of genes in the ranked list before the maximum ES is reached. "Tags" refers to the percentage of gene hits that are in the leading edge. "List" is the percentage of genes in the dataset that are in the leading edge. Signal is a combination of the previous two numbers, where a higher number indicates that the genes in the geneset are clustered closer towards the top of the ranked dataset.

E2

| Gene | Protein_Change | VCF Phred | Zygosity | Alternate reads | Total reads | VAF | Chr | Position | Ref_Base | Alt_Base | UniProt Accession | CHASMplus Score | CHASMplus_ GBM Score |
|---|---|---|---|---|---|---|---|---|---|---|---|---|---|
| EGFR | p.Ala289Val | 617 | het | 25 | 36 | 0.69 | chr7 | 55154129 | C | T | P00533 | 0.80 | 0.85 |
| TP53 | p.Arg273His | 13568 | het | 504 | 587 | 0.86 | chr17 | 7673802 | C | T | P04637 | 0.82 | 0.75 |
| MAP3K1 | p.Asp806Asn | 2134 | het | 91 | 234 | 0.39 | chr5 | 56881616 | G | A | Q13233 | 0.62 | 0.32 |
| MAP3K1 | p.Val906Ile | 2833 | het | 113 | 251 | 0.45 | chr5 | 56881916 | G | A | Q13233 | 0.35 | 0.18 |
| USP9X | p.Gln640His | 731 | hom | 23 | 23 | 1 | chrX | 41162812 | A | C | Q93008 | 0.52 | 0.18 |
| PTCH1 | p.Pro1315Leu | 563 | het | 26 | 57 | 0.46 | chr9 | 95447312 | G | A | Q13635 | 0.33 | 0.17 |
| LZTR1 | p.Gly301Ser | 4774 | hom | 161 | 161 | 1 | chr22 | 20991737 | G | A | Q8N653 | 0.34 | 0.16 |
| APC | p.Val1822Asp | 1889 | hom | 61 | 61 | 1 | chr5 | 112841059 | T | A | P25054 | 0.49 | 0.16 |
| PIK3R2 | p.Ser234Arg | 13717 | hom | 437 | 437 | 1 | chr19 | 18161380 | A | C | O00459 | 0.35 | 0.13 |
| CDKN2A | Aligned RNAseq reads completely absent from CDKN2A locus on chr 9. | | | | | | | | | | | | |

G7

| Gene | Protein_Change | VCF Phred | Zygosity | Alternate reads | Total reads | VAF | Chr | Position | Ref_Base | Alt_Base | UniProt Accession | CHASMplus Score | CHASMplus_ GBM Score |
|---|---|---|---|---|---|---|---|---|---|---|---|---|---|
| TP53 | p.Arg248Gln | 7489 | het | 292 | 603 | 0.48 | chr17 | 7674220 | C | T | P04637 | 0.85 | 0.80 |
| TP53 | p.Arg282Trp | 10846 | het | 431 | 902 | 0.48 | chr17 | 7673776 | G | A | P04637 | 0.79 | 0.70 |
| MAP3K1 | p.Asp806Asn | 264 | het | 11 | 17 | 0.65 | chr5 | 56881616 | G | A | Q13233 | 0.62 | 0.32 |
| SETD2 | p.Pro1962Leu | 9043 | hom | 303 | 303 | 1.00 | chr3 | 47083895 | G | A | Q9BYW2 | 0.32 | 0.21 |
| MAP3K1 | p.Val906Ile | 528 | het | 21 | 35 | 0.60 | chr5 | 56881916 | G | A | Q13233 | 0.35 | 0.18 |
| APC | p.Val1822Asp | 556 | het | 25 | 49 | 0.51 | chr5 | 112841059 | T | A | P25054 | 0.49 | 0.16 |
| PIK3R2 | p.Ala415Thr | 12598 | het | 489 | 1026 | 0.48 | chr19 | 18163100 | G | A | O00459 | 0.40 | 0.14 |
| CDKN1A | p.Asp149His | 39 | het | 5 | 18 | 0.28 | chr6 | 36684546 | G | C | P38936 | 0.59 | 0.13 |
| PIK3R2 | p.Ser234Arg | 14405 | hom | 464 | 464 | 1.00 | chr19 | 18161380 | A | C | O00459 | 0.35 | 0.13 |
| ARID1A | p.Pro683Ser | 1407 | het | 65 | 196 | 0.33 | chr1 | 26760982 | C | T | O14497 | 0.67 | 0.12 |

**Supplementary table 5. Potential GBM cancer driver mutations in G7 and E2 cell lines**

The table shows the missense mutations in G7 and E2 cells that are predicted to have cancer driver activity in glioblastoma.

DNA sequence changes in VCF files were generated from aligned RNAseq data from control (untreated) cells compared to the Hg38 reference sequence. Open-Cravat was used to identify missense mutations that have a CHASMplus p value < 0.01 (i.e. predicted to be a cancer driver mutation) and CHASMplus GBM p value < 0.05 (i.e. predicted to be a glioblastoma diver mutation). CHASMplus uses machine learning to discriminate between cancer driver or passenger mutations (Tokheim and Karchin, 2019). The CHASM score is a value between 0 and 1 that indicates the predicted cancer driver activity. The higher the score, the more powerful the driver mutation.

Zyg: zygosity. Alt reads: number of reads matching the alternate allele. VAF: variant allele frequency. Chr: chromosome. Ref: reference base. Alt: alternate base

IGV software was used to show that no reads aligning to the CDKN2A locus were detected in E2 cells, and to infer that the G7 TP53 variants chr17:7674220 (GRCh38) and chr17:7673776 (GRCh38) are located on different DNA alleles. Loss of both CDKN2A alleles was confirmed by alignment of reads from genomic DNA from the ChIP-seq samples.

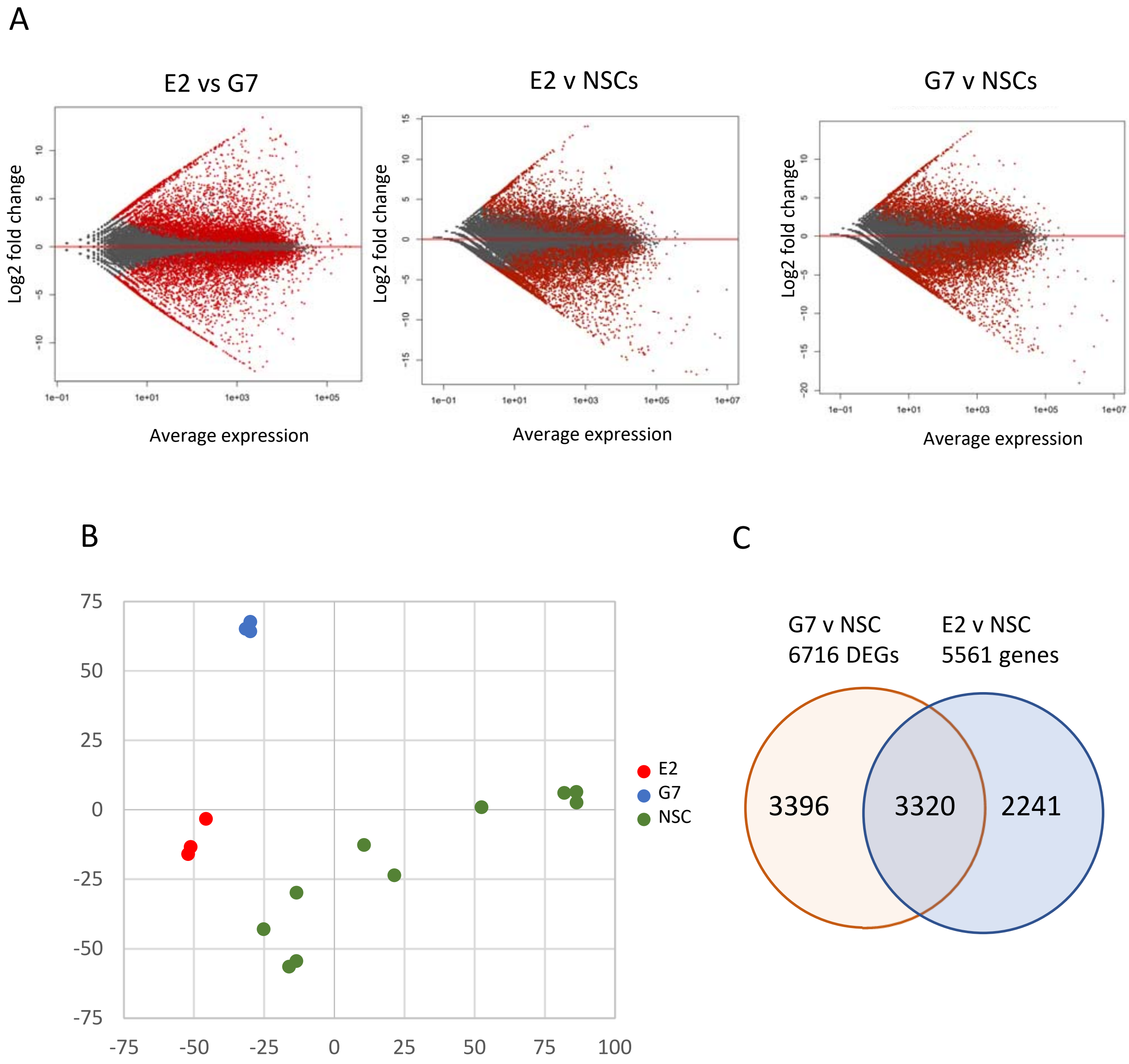


**Supplementary figure 1:** Characterisation of GBM cell lines through RNAseq analysis

A) MA plots showing comparison of RNAseq data from the indicated samples. X axis: mean of the normalised gene expression counts from all samples included in the analysis. Y axis: log2 fold change of normalised gene expression. Red spots: adj.p < 0.1
B) PCA plot showing the relationship between E2, G7 and normal NSC lines.
C) Venn diagram of differentially expressed genes (adjp < 0.01, log2fc > 1 or < -1) in GBM cell lines compared to normal NSCs.

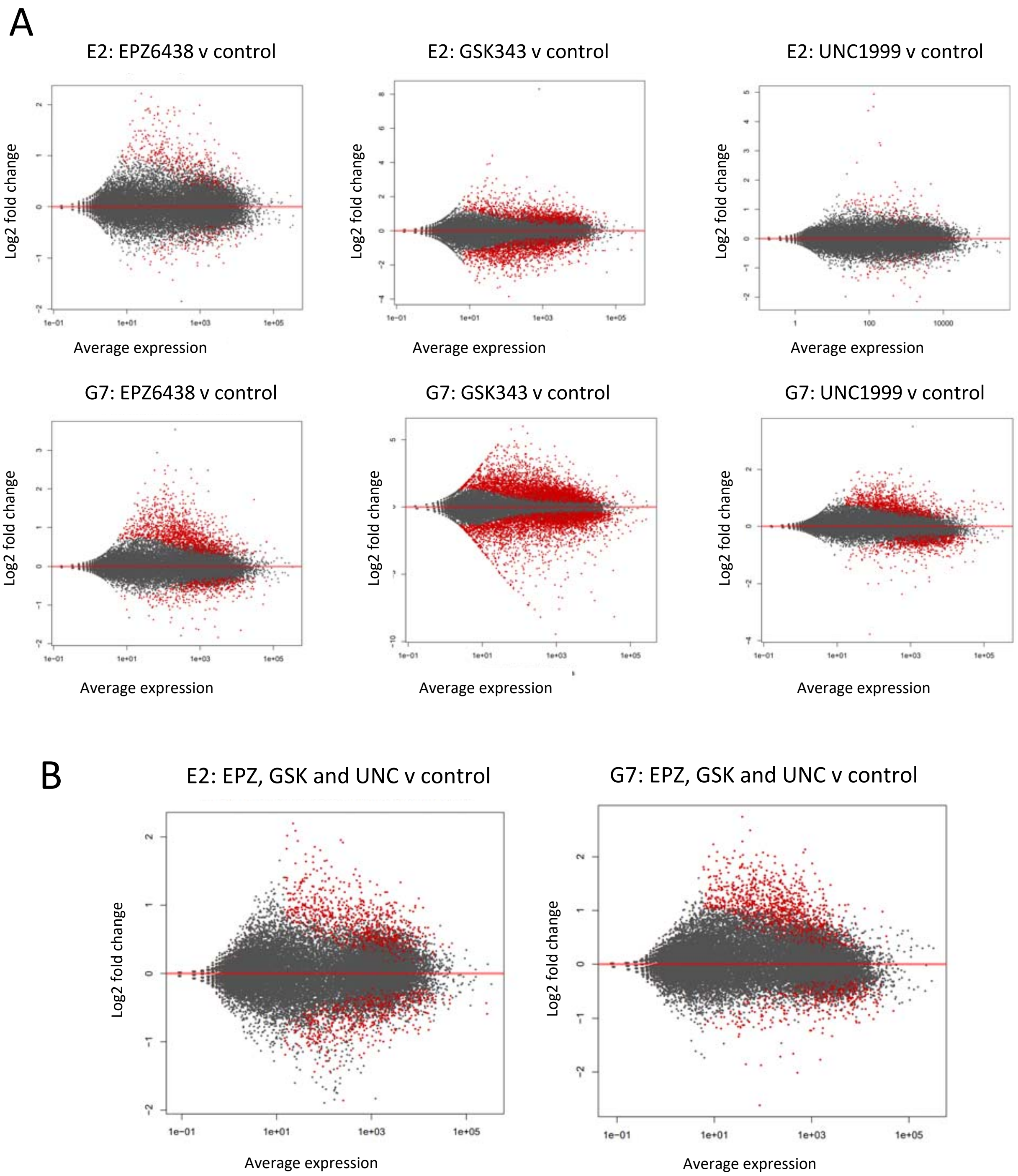


**Supplementary figure 2: MA plots from DESEQ2 analyses on the indicated datasets.**

A: pairwise analyses of RNAseq data from drug treated cells versus DMSO control
B: group analyses of RNAseq data from all drug treated cells versus DMSO control
Cells were treated with 2 µM EZH2 inhibitor for 5 days prior to harvesting and paired end RNA sequencing, in triplicate.
X axis: mean of the normalised gene expression counts from all samples included in the analysis.
Y axis: log2 fold change of normalised gene expression in experimental versus control. Red spots: adj.p < 0.1

.

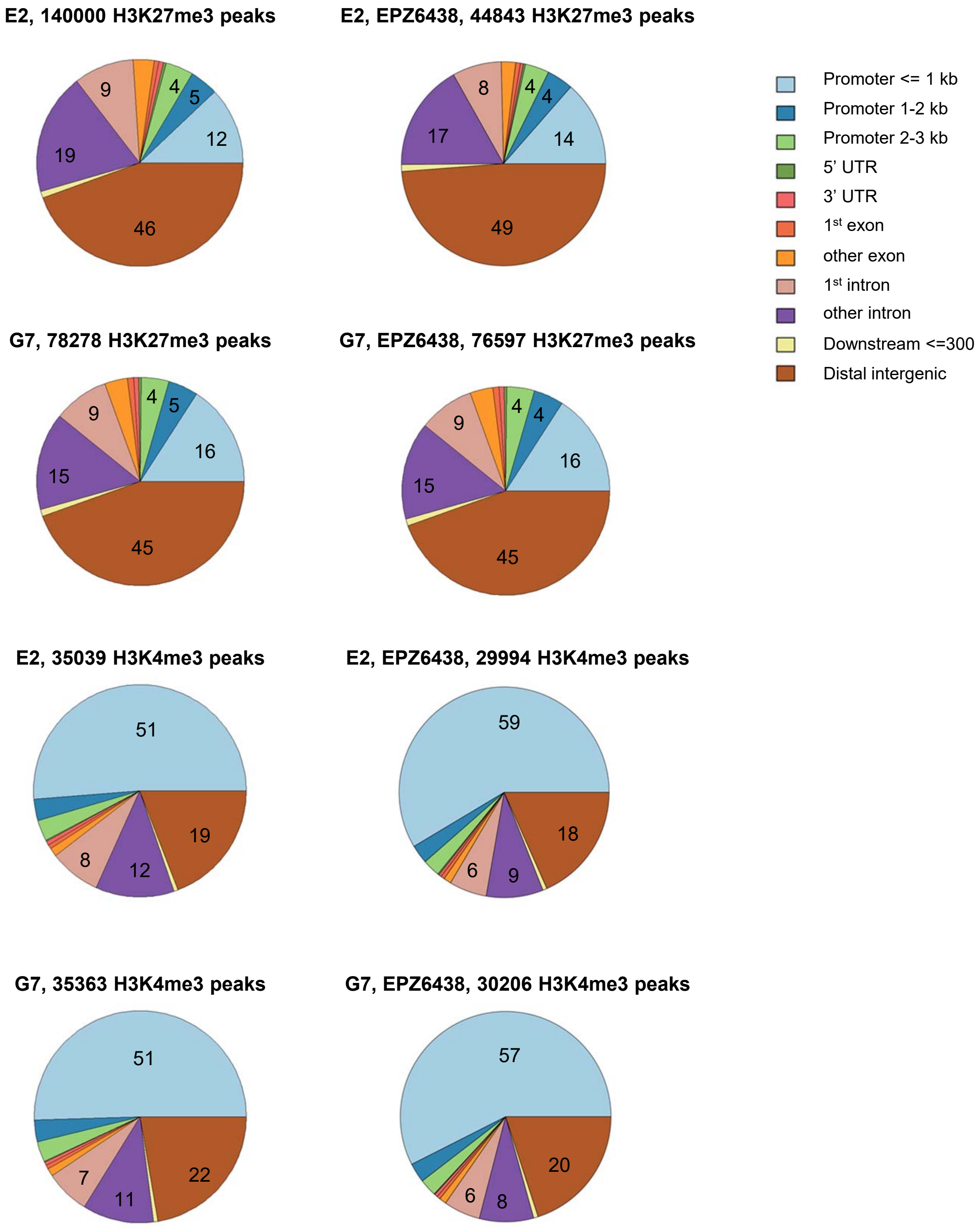


**Supplementary figure 3: Distribution of H3K27me3 and H3K4me3 peaks in E2 and G7 cells with respect to genetic elements.**

ChIP-Seq for H3K27me3 and H3K4me3 was performed in E2 and G7 cells treated with 2 μM EPZ6438 for 5 days, or DMSO control. Reads were aligned to Hg38, peaks were called using MACS2 broad peak by comparison to input control. Overlaps of peaks with gene regions was calculated using Chipseeker. The percentage of peaks in the largest categories are indicated.
